# Neural manifold connectomics reveals multiregime functional connectivity

**DOI:** 10.64898/2026.07.24.740595

**Authors:** Puneet Velidi, Enrico Amico, Farouk Nathoo, Michelle F Miranda

## Abstract

Neural activity is organized in low-dimensional structure, yet functional connectivity in fMRI typically represents each brain parcel by a single voxel-averaged time series. This scalar representation makes whole-brain connectivity tractable but discards potentially informative dimensions of within-parcel BOLD activity. Here, we represent each parcel by a low-dimensional temporal subspace derived from its principal-component time series and use the RV coefficient to quantify connectivity between regional subspaces. Across Human Connectome Project resting-state and working-memory data, progressively expanding these subspaces reveals reproducible connectivity regimes with distinct network and identifiability profiles. At rest, connectivity constructed from the first principal component identifies individuals more strongly than either voxel-averaged functional connectivity or higher-dimensional subspace representations. During working memory, identifiability instead peaks after secondary components are included, indicating that the distribution of individual-specific information across the regional PCA spectrum depends on cognitive state. These patterns replicate across independent samples and remain robust across multiple parcellation resolutions. Together, our findings show that within-parcel BOLD structure contains identity- and statedependent information that is obscured by scalar regional summaries. Functional connectivity may therefore be better understood as a family of related connectomes indexed by the regional subspace retained, providing a general framework for mapping interactions between low-dimensional neural representations.

## 1 Introduction

Neural activity, across scales from single-neuron populations to whole-brain Blood Oxygen Level Dependent (BOLD) activity, consistently exhibits low-dimensional structure reflecting the redundancy of biological neural systems [1]. This structure is often described as neural manifolds, low-dimensional structures in neural state space along which population activity is constrained to evolve, shaped by biophysical constraints such as the behavioral demands placed on neural circuits [2, 3]. In systems neuroscience, neural manifolds have reframed decades of single-neuron findings into a population-level framework, with a small number of neural modes serving as the basic building blocks of neural dynamics rather than the activity of individual units [2].

Functional connectivity (FC) in fMRI summarizes temporal dependencies between brain regions, and group- and individual-level differences in FC are used to characterize many aspects of the brain–behavior relationship. Pearson’s (r) FC, which we refer to as ‘standard” FC throughout this paper, has become a cornerstone of human neuroimaging, in part due to its ability to uniquely identify individuals. FC profiles act as ‘fingerprints” that reliably distinguish one person from another across scanning sessions, task conditions, and imaging sites [4, 5]. Connectome identifiability [6] provides insights into between-individual variability in network organization, motivating its use in precision neuroscience and personalized clinical assessment [7–9]. Disease may alter both the strength and anatomical composition of these fingerprints. For instance, in MEG, healthy control brains have shown greater identifiability than brains of patients with neurological disease [10], while individual FC profiles remain identifiable in mild cognitive impairment and Alzheimer’s disease even as the connections supporting identification are reconfigured with disease progression [11]; in Parkinson’s disease, patient-specific fingerprint features have additionally been associated with motor severity and cognitive performance and shown to distinguish patients with and without minor hallucinations [12].

We argue that standard functional connectivity compresses rich within-region dynamics into a single representative signal. While this simplification has been effective for studying large-scale brain organization, it also raises the possibility that important aspects of neural organization remain hidden in dimensions of the BOLD signal that are routinely discarded. At the neuronal scale, low-dimensional population manifolds emerge in cortical and subcortical circuits despite sparse connectivity [1, 2]; at the whole-brain scale, principal component analysis (PCA) of parcel-averaged BOLD recovers a cognitive manifold spanning multiple tasks [13]. Relevant to our analysis, at the voxel-to-ROI scale, the latent dynamics of BOLD have not been systematically related to functional connectivity. If within-parcel BOLD can be summarized by a small number of principal components, then averaging accesses only one direction of that structure, and neither the remaining variance nor its relationship to fingerprinting capacity has been systematically investigated. Here, we are particularly interested in fingerprinting as it serves as a basic test of our method’s validity for population level inference.

Motivated by this gap, we ask whether low-dimensional structure within parcels provides a richer account of functional connectivity than conventional ROI averaging, and whether this structure contributes to individual identity and network organization across brain states. To quantify relationships between parcel-level principal-component time series, we use the RV coefficient, a multivariate generalization of Pearson correlation that measures association between sets of time series rather than between individual signals [14]. This formulation makes the amount of within-region variance retained an explicit modeling choice. Prior work has generally fixed this quantity when estimating RV-based connectivity [15, 16], leaving unclear how different portions of within-region structure contribute to functional connectivity, network organization, and brain fingerprinting across cognitive states.

We therefore systematically vary the amount of within-region structure retained in each regional representation and examine how connectivity, network organization, and identifiability change as additional latent dimensions of BOLD activity are incorporated. Within-ROI BOLD variance is concentrated in a relatively small number of principal components, extending to the voxel-to-ROI scale the redundancy observed at neuronal and whole-brain scales [1, 13]. At rest, differential identifiability is highest for connectivity constructed from the first principal component and then declines monotonically as additional components are retained, while remaining higher than that of Pearson FC at low variance-explained thresholds, including RV_10_ and RV_20_. This result indicates that parcel averaging is not necessarily the optimal regional representation for preserving individual-specific connectivity information.

We further find that the PCA spectrum separates into functionally distinct bands whose connectivity profiles differ in network organization, identifiability, and similarity to other bands (Fig. 1). The distribution of individual-specific information across these bands is state-dependent: whereas resting-state identifiability is strongest in the dominant component, working-memory identifiability peaks when components collectively explaining approximately 20–60% of within-ROI variance are retained. Together, these findings indicate that within-parcel BOLD contains structured, functionally meaningful information beyond the parcel average, and that different portions of this structure carry individual-specific information across cognitive states. Functional connectivity may therefore be better understood as a family of related matrices indexed by the regional subspace retained, rather than as a single matrix determined by one regional summary. We replicate these findings in independent Human Connectome Project samples and show that they are robust across multiple parcellation resolutions [17, 18].

**Fig. 1:**
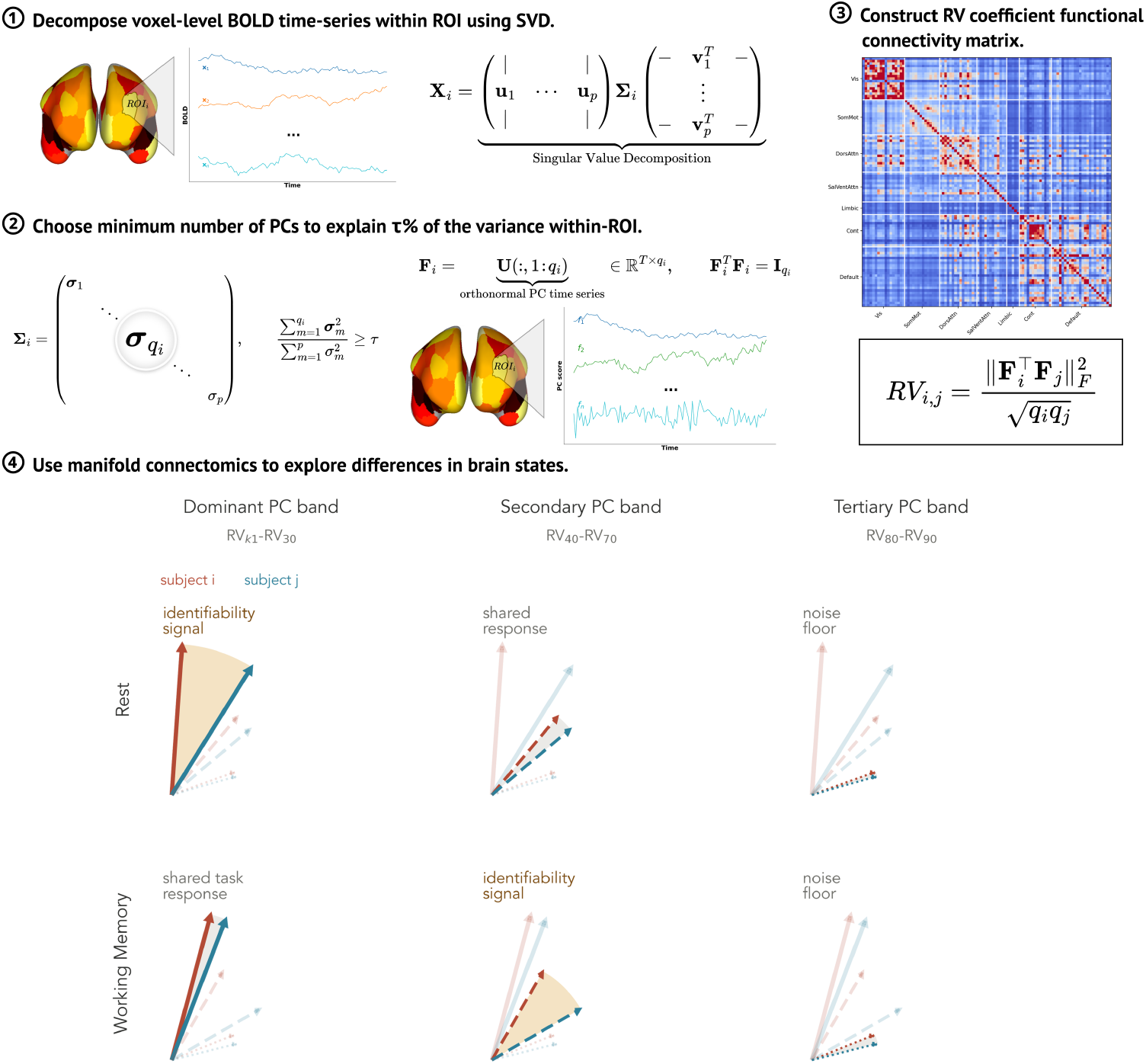
Overview of the RV-based functional connectivity computation for a given variance threshold *τ* and hypothesized PC band. Arrows denote subject-specific PC modes within an ROI, with arrow length schematically indicating variance explained. Angular separation between the two subjects’ arrows within a band represents between-subject divergence in that mode, interpreted as greater individual-specific or identifiability signal. Small angular separation indicates a more shared response, whereas the tertiary band is shown as noise-dominated.

## 2 Results

### 2.1 Leading-component RV mirrors scalar FC at rest

To assess how multivariate representations relate to standard functional connectivity, we compared RV-based connectivity matrices across variance thresholds to Pearson correlation-based FC. At rest, the group FC matrix of RV_*k*1_ closely resembles Pearson R (*r* = 0.92; Fig. 2A). This similarity decreases monotonically as additional PCs are incorporated: *r* = 0.88 at RV_10_, *r* = 0.71 at RV_20_, continuing through *r* = 0.57, 0.48, 0.43 at RV_30_–RV_50_ (Supplementary Fig. S1), and falling to *r* = 0.35, 0.23, 0.14, 0.08 at RV_60_–RV_90_ (Supplementary Fig. S2). Initially, as the correlation with R diminishes and more principal components are included in parcel subspaces, betweennetwork manifold-based connectivity becomes increasingly sparse and within-network connectivity sharpens i.e the blocks on the diagonal of the matrices Fig. S1S2 which we elaborate on in Figure 5. Then, at RV_60_ and above, the connectivity matrices approach a homogeneous, disconnected graph with no discernible network structure.

**Fig. 2:**
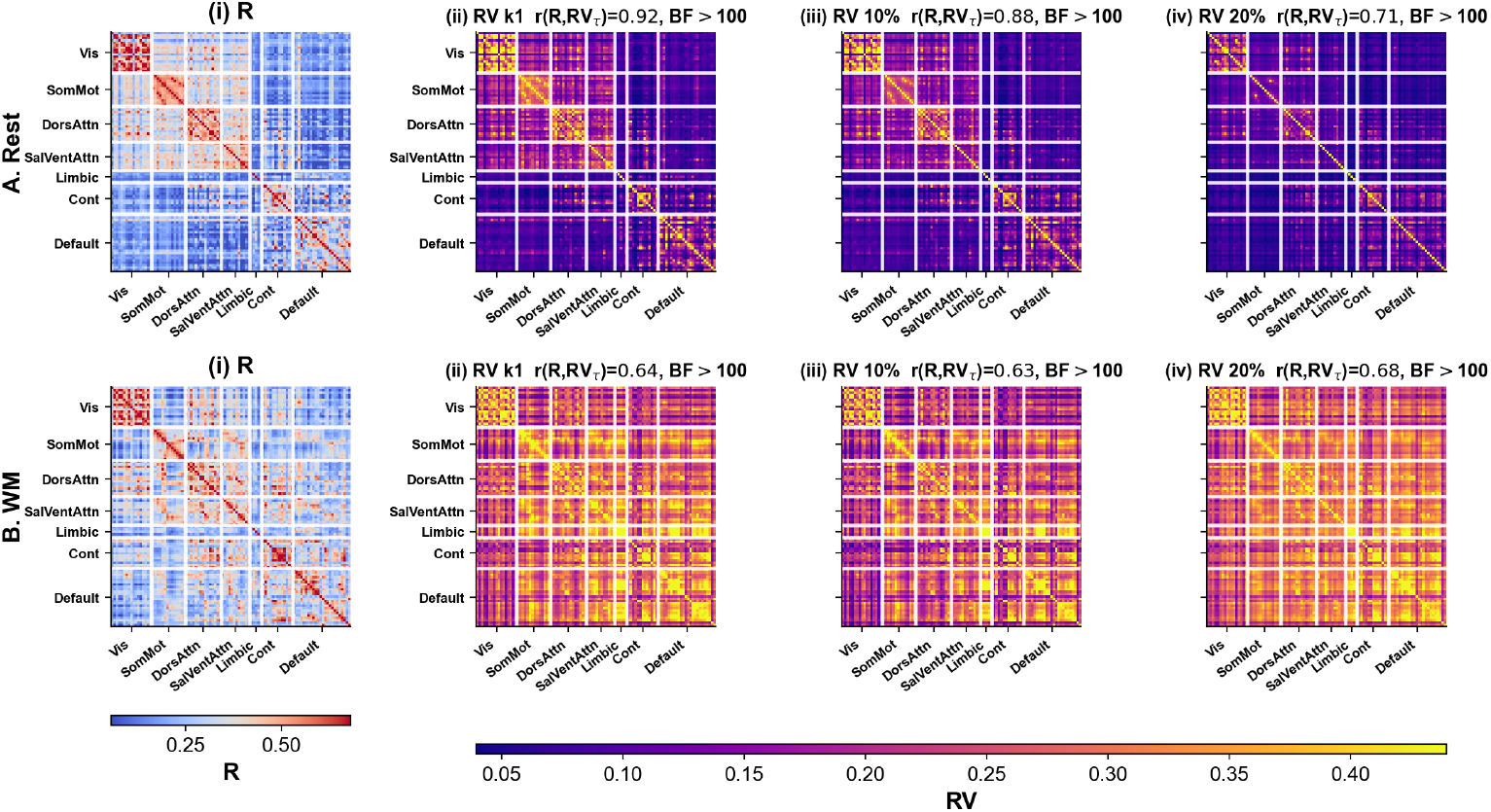
Group FC matrices for Pearson correlation (R) and RV-based FC (RV) across variance thresholds. A., group resting-state FC matrices, averaged across REST1 and REST2 scans and all confirmatory subjects; B., group WM task FC matrices averaged across phase encoding directions and all subjects. i. Pearson’s R-based FC. RV-based FC matrices retaining ii. the first PC as well as the PCs that explain at least iii. 10% and iv. 20% of the within-ROI variance. Bayes factors in the figure are one-sided default Bayes factors (BF_10_) for *H*_1_ : *r >* 0. All comparisons showed decisive evidence for a positive correlation (BF_10_ *>* 100).

During WM, the pattern differs. The RV–R correlation does not peak at the first PC but at RV_20_ (*r* = 0.68), and the decline with increasing threshold is shallower (*r* = 0.64 at RV_30_, *r* = 0.53 at RV_40_, *r* = 0.47 at RV_50_, reaching *r* = 0.44 at RV_60_ and *r* = 0.24 at RV_90_). While R FC shows generally lower connectivity magnitudes during WM than at rest, this pattern reverses in RV: WM RV FC matrices are denser and maintain stronger between-network connectivity across the full spectrum of *τ* . Even at RV_80_–RV_90_, WM RV FC retains visible between-network structure, which is likely due to task recruitment of integrative cross-network cognitive processes. Rest R FC has high similarity to some RV matrices, but task R FC does not closely resemble any RV matrix.

An alternative decomposition constructs FC matrices from only the PCs that explain the additional variance between consecutive thresholds rather than the cumulative subspace. These band-specific RV matrices are shown in Supplementary Figs. S10–S13.

### 2.2 Within-ROI variance spans multiple PCs

We examined the eigendecomposition of the within-ROI data matrix to validate that changing the variance explained threshold is, in fact, meaningful - if a single component explained nearly all within-ROI variance, then the RV framework would collapse to a one-dimensional representation. Instead, the number of PCs per parcel needed to explain *τ* % variance increases non-linearly with *τ* and shows greater spread at higher thresholds (Fig. 3B). This is consistent with a power law singular value spectrum, where the first few components explain most of the variance and later components provide, on average, additional variance explanation, albeit with decreasing consistency in doing so (Fig. 3C). The mean number of PCs needed to explain within-ROI variance also varies non-trivially across the brain, reflecting both differences in ROI size and differences in regional activity complexity (Fig. 3A). Since PCA is bounded by the rank of the centered data matrix, the maximum number of principal components per ROI is min(TRs − 1, voxels), which is 1199 for resting state and 404 for WM. In this dataset, WM generally requires fewer PCs, consistent with stronger task-constrained manifold structure during the *N* -back acquisition.

**Fig. 3:**
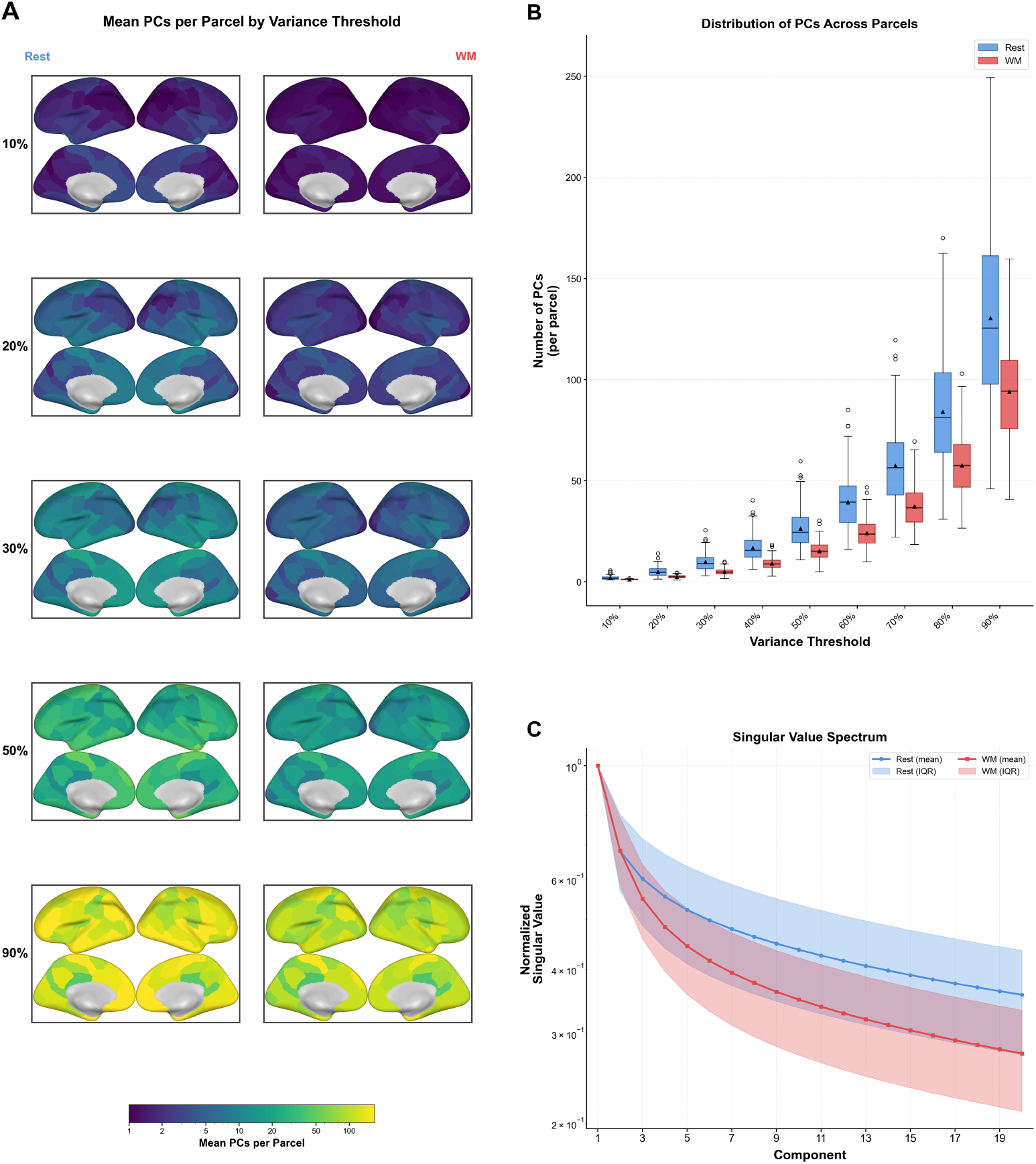
PCs needed to explain different levels of variance in resting-state and WM BOLD ROI time series. A., mean number of PCs across subjects to explain (10%, 20%, 30%, 50%, 90%) of the within-ROI variance in the Schaefer 100 parcellation in the confirmatory sample. B., box plots showing the distribution of PCs across all subjects and all ROIs to explain (10%, 20%, 30%, 50%, 60%, 70%, 80%, 90%) of the within-ROI variance. C., distribution of singular values across subjects and ROIs (shown by mean and IQR) as a function of PC number.

### 2.3 RV reveals three identifiability regimes

We next used between-threshold similarity to ask whether different portions of the PC spectrum carry similar or distinct connectivity information. RV-based FC revealed distinct regimes of PC activity: RV_*k*1_–RV_30_ matrices are similar to each other but dissimilar to those at other thresholds, with a similar block structure for the RV_40_–RV_70_ and RV_80_–RV_90_ regimes (Fig. 4A). For both conditions, the means of the distributions of *I*_self_[6](the test–retest stability of a single individual’s scans, see methods) and *I*_others_ (the test–retest stability between any two individuals’ scans, see methods), increase monotonically with *τ*, with the distributions themselves becoming more compressed (Fig. 4B & D). Our primary finding is that, in resting state, as additional within-parcel variance is incorporated, fingerprinting performance peaks at the first principal component and then monotonically decreases, still outidentifying Pearson’s R-based FC at lower variance explained thresholds (i.e., RV_10_, RV_20_; Fig. 4E). In WM, almost all RV FC matrices are more identifiable than R FC, with a broad intermediate-dimensional peak across RV_30_–RV_60_ (Fig. 4E). For both resting state and working memory, the *I*_diff_ profiles were highly concordant across exploratory and confirmatory samples (rest: *r* = 0.985, BF_10_ = 4.6 *×* 10^4^; WM: *r* = 0.953, BF_10_ = 1.6 *×* 10^3^).

**Fig. 4:**
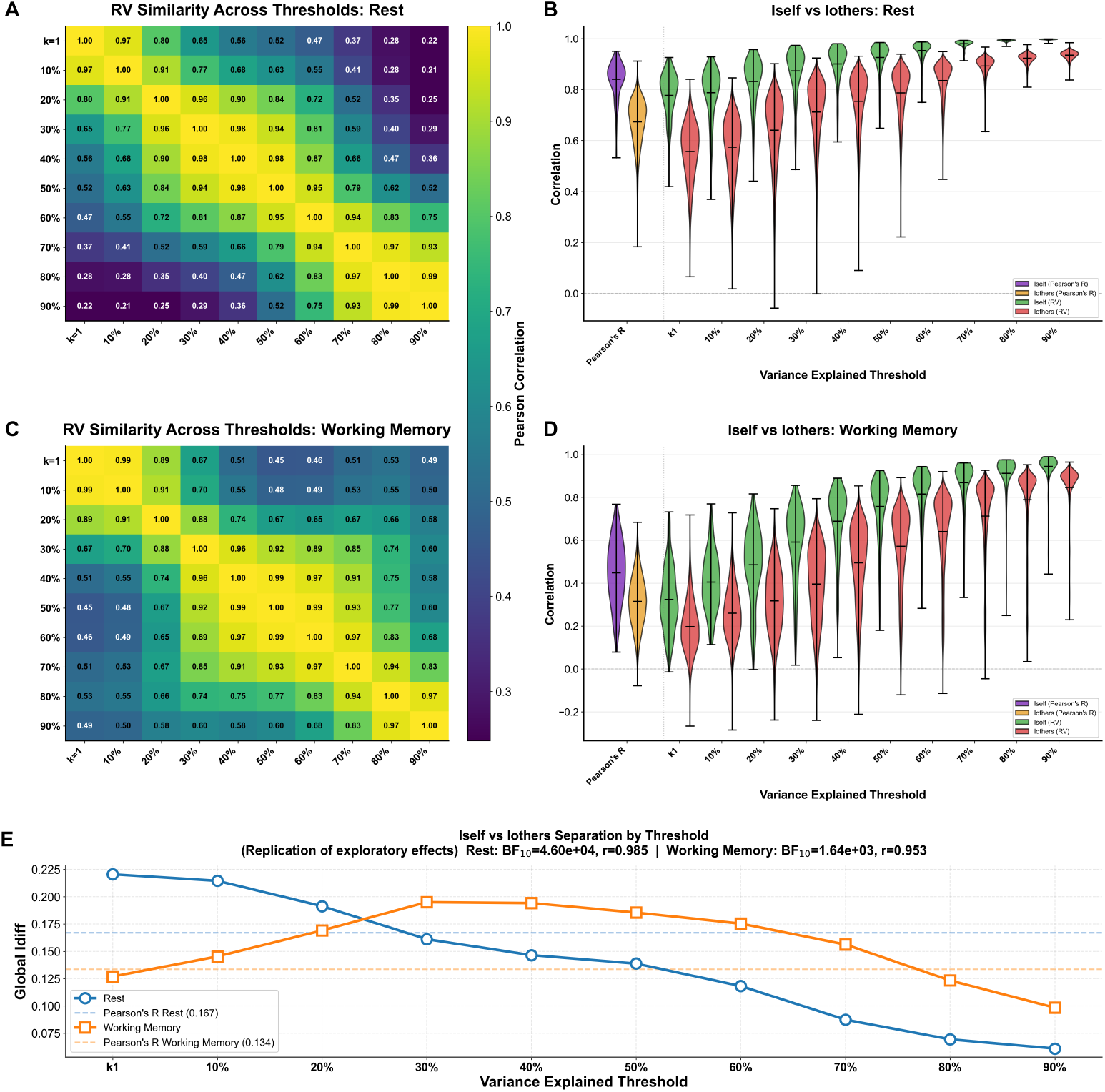
Between-threshold RV-based FC similarity and identifiability. A. Between-threshold resting-state RV-based FC similarity (*r*) from RV_*k*1_ to RV_90_, averaged across subjects. B. Violin plots of the distribution of each individual’s identifiability with themselves across test and retest (*I*_self_, green) and with other subjects’ retest (*I*_others_, red) for rest RV-based FC at each threshold. C. Between-threshold WM RV-based FC similarity (*r*) from RV_*k*1_ to RV_90_, averaged across subjects. D. Violin plots of *I*_self_ (green) and *I*_others_ (red) for WM RV-based FC at each threshold. E. Differential identifiability (*I*_diff_ = *I*_self_ *− I*_others_) across thresholds (blue: resting state; orange: WM). Dotted lines indicate *I*_diff_ of Pearson’s R-based FC for resting state (0.167) and WM (0.134). For replication of exploratory effects, the exploratory and confirmatory *I*_diff_ values were correlated across variance thresholds (see Methods). Rest: *r* = 0.985, BF_10_ = 4.6 ×10^4^; WM: *r* = 0.953, BF_10_ = 1.6 × 10^3^ (one-sided correlation BF, *H*_1_: *ρ >* 0 vs *H*_0_: *ρ* = 0; *n* = 10 thresholds), indicating decisive evidence for replication in both conditions.

### 2.4 Low-dimensional RV segregates rest and integrates WM

We used Yeo 7 network radar plots to interpret the RV matrices at the level of canonical functional networks. RV_20_ shows segregation of networks at rest, with high within-network connectivity but low between-network connectivity (Fig. 5A(i)). The same threshold shows integration of networks in WM, with both high within-network and high between-network connectivity (Fig. 5B). At rest, RV_20_ shows the greatest within-network connectivity in the visual network, whereas RV_80_ shows the greatest within-network and between-network connectivity in the limbic network. In WM, the overall within- and between-network connectivity (the areas of the radar polygons) generally decreases with variance explained. At rest, these areas are largest at early PCs, then contract at intermediate PCs, and finally expand slightly within-network and to a greater extent between-network (Supplementary Fig. S3). In Figure 5A and Supplementary Fig. S3, at rest, we observe greater within-limbic-network connectivity compared to other networks at high variance explained thresholds (Supplementary Fig. S3A(iii)) and lesser within-limbic-network connectivity compared to other networks at low variance explained thresholds (Supplementary Fig. S3A(i)).

**Fig. 5:**
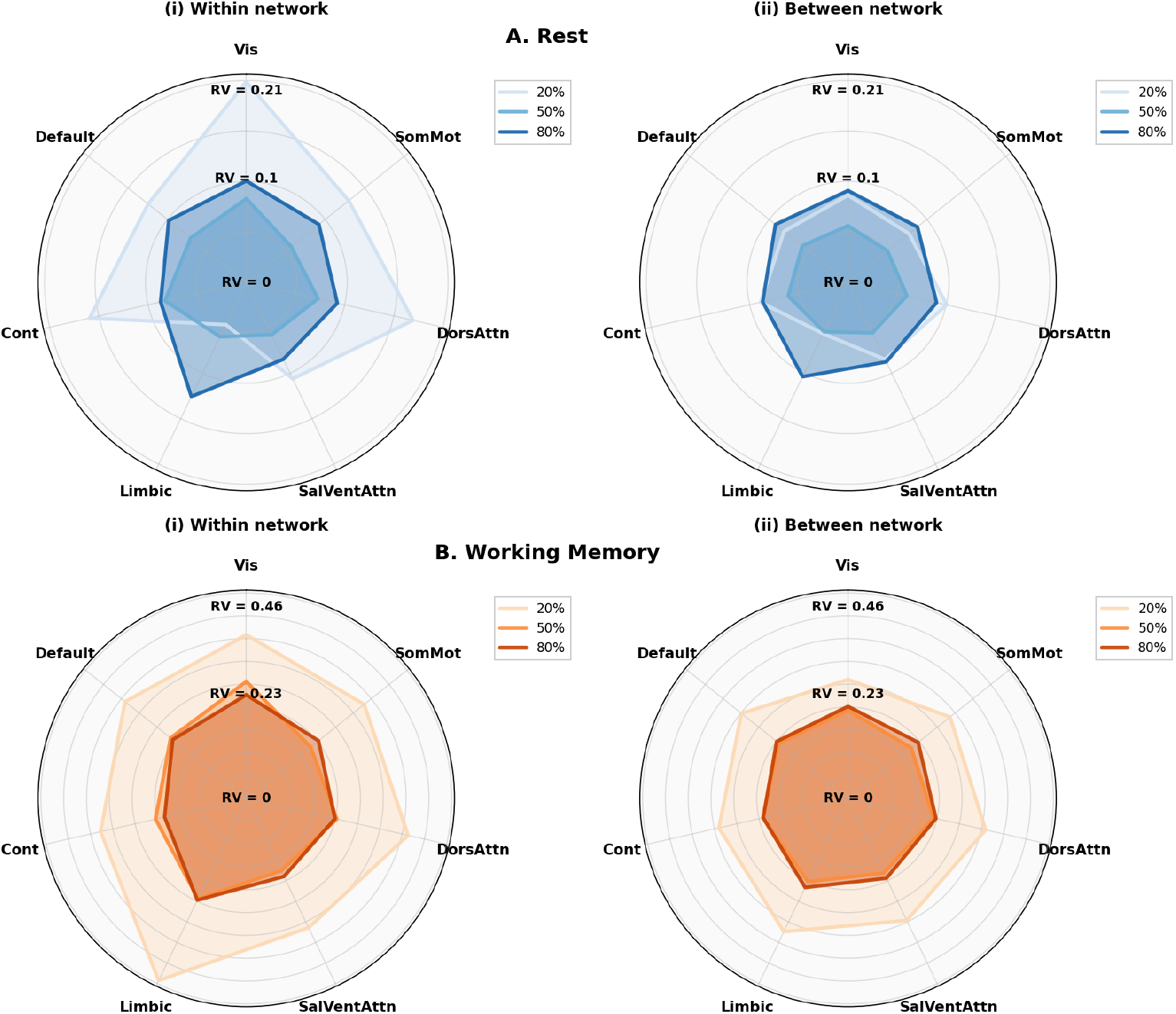
A. Yeo 7 network-level radar plots for group resting-state RV-based FC in the confirmatory sample; B. Yeo 7 network-level radar plots for group WM RV-based FC in the confirmatory sample. i. Within-network: mean connectivity among ROIs within a given network. ii. Between-network: mean connectivity of ROIs within a given network with ROIs outside the network. RV_20_, RV_50_, and RV_80_ were taken as representative of low, moderate, and high variance explained FCs. See the full set of radar plots in the Supplementary Information. Contours darken with increasing variance explained threshold.

### 2.5 Parcel identifiability declines with increase in variance-explained

We next decomposed identifiability by parcel (see Methods 4) to examine which regional connectivity profiles carry individual-specific information. In resting state, parcels become less identifiable with increasing variance explained (Fig. 6A & Supplementary Fig. S4(i)), with a steep drop after *τ* = 20%, after which *I*_diff_ approaches 0. In WM, parcels become more identifiable until *τ* = 20% and remain, in absolute terms, identifiable past this point. In both WM and rest, all parcels show low identifiability at *τ* = 80% (Fig. 6A(iii) and B(iii)). The rank ordering of parcel identifiability shows approximate self-similarity (Fig. 6A(iv) and B(iv)), with parcel-level transition zones shifted relative to the global matrix regimes: low thresholds (RV_*k*1_–RV_10_), intermediate thresholds (RV_20_–RV_70_), and high thresholds (RV_80_–RV_90_). In Fig. 6B(iv), the second regime is not self-similar until *τ* = 40%; that is, RV_20_ and RV_30_ fall outside the block-diagonal structure. Overall, there is greater rank stability between WM RV FCs. Notably, in rest (Fig. 6A(iv)), the parcel rank orderings become anti-correlated between high and low variance explained thresholds, suggesting that different portions of the PCA spectrum have distinct spatially-varying fingerprints that may be independent of the gradient we observe of fingerprinting decreasing with variance explained.

**Fig. 6:**
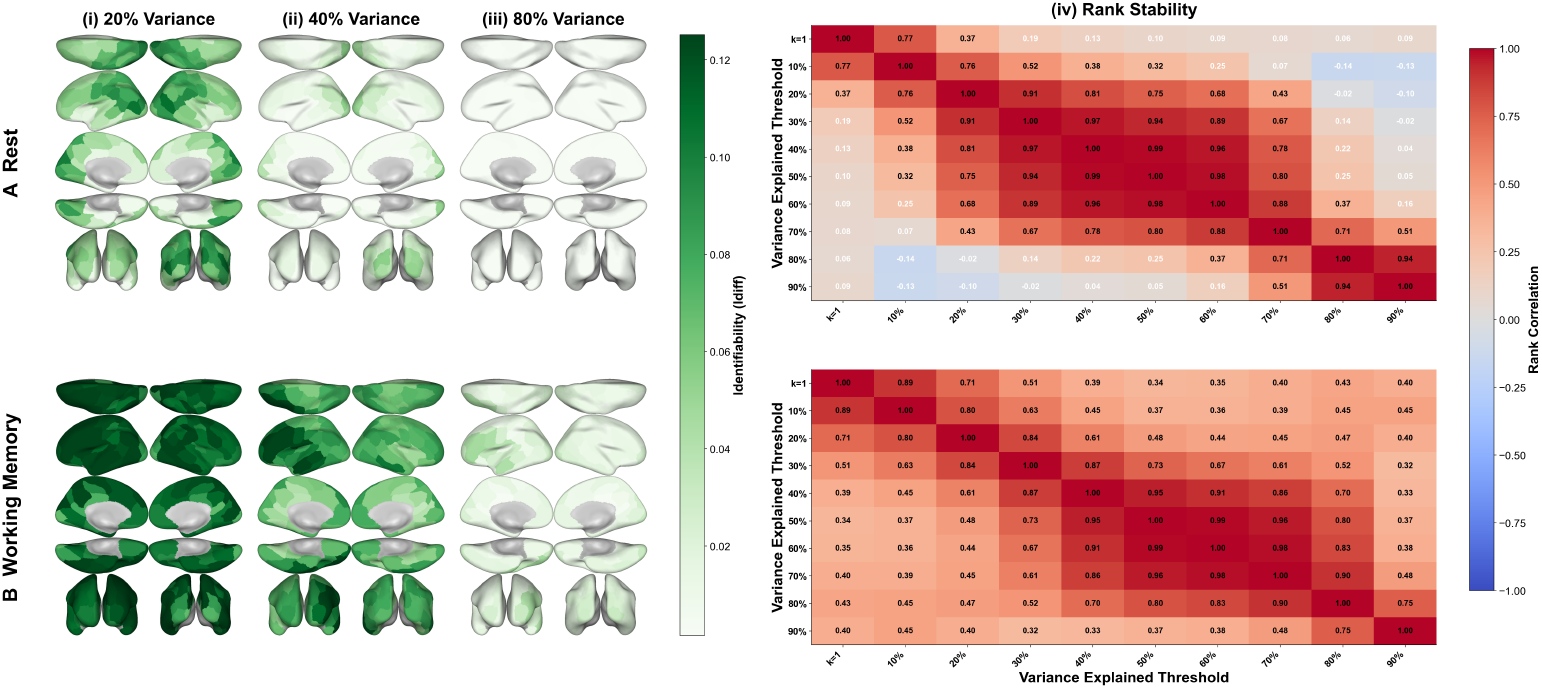
Parcel-level identifiability and ranking stability. A. Resting-state-derived parcel identifiability; B. WM-derived parcel identifiability. i. 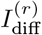 from RV_20_ FC; ii. 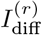 from RV_40_ FC; iii. 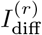 from RV_80_ FC; iv. rank stability of parcel identifiability rankings across variance thresholds.

## 3 Discussion

The discussion develops three implications of the RV framework. First, it reframes functional connectivity as dependence between multivariate regional representations, making the choice of parcel representation an explicit part of the connectome and clarifying how subspace FC relates to conventional scalar FC. Second, varying the retained PCA structure reveals a hierarchical organization of dominant, intermediate, and later regional modes with distinct network and identifiability properties. Third, because RV measures alignment between sets of latent time series rather than relying specifically on PCA, it provides a general framework for defining connectivity between low-dimensional representations across spatial scales, cognitive states, and potentially imaging modalities.

### 3.1 Scalar and manifold connectomes

Pearson’s R FC remains a useful reference point because it defines the dominant scalar connectome used throughout human neuroimaging. Its success in brain fingerprinting shows that a single regional summary can preserve stable individual differences [4]. However, parcel averaging also makes a strong assumption that the relevant activity of each region is well represented by one time series. This assumption is convenient, but it is not optimal in that it removes many degrees of freedom in the BOLD signal. Within-parcel heterogeneity is well documented [19, 20], task-evoked responses can vary substantially across voxels within the same region [21], and parcellation choices can distort clinical group comparisons [22]. In this setting, RV FC is a way to make explicit which dimensions of the within-parcel PCA decomposition are being used to construct a connectome.

The relationship between R and RV depends strongly on cognitive state. At rest, RV_*k*1_ closely resembles Pearson FC (Fig. 2A), yet it fingerprints individuals better than R (Fig. 4E). This indicates that small differences between the parcel average and the dominant within-parcel component contain reliable identity information. The parcel-level pattern, with strong low-threshold identifiability in frontoparietal and default mode regions (Supplementary Fig. S4A(ii)), is consistent with prior fingerprinting work [4]. As additional PCs are added at rest, RV matrices become progressively less R-like and less identifiable, suggesting that the dominant component of the resting PCA decomposition carries most stable individual information while higher components add increasingly shared or unstable variance.

Working memory shows the opposite pattern. R and RV are much less similar during WM than at rest (Fig. 2B), and RV identifiability peaks at intermediate thresholds (Fig. 4E). This suggests that task fMRI does not necessarily suppress individual differences; rather, those differences may be expressed in different portions of the PCA spectrum. One interpretation is that task structure aligns the leading component across individuals while pushing subject-specific implementation into secondary components. These secondary components may reflect differences in attention, strategy, processing efficiency, or local response heterogeneity during the *N* -back task. Because many common parcellations, including Schaefer, are derived from resting-state organization [23], scalar task FC may be especially likely to average over task-induced within-parcel structure.

In both conditions, *I*_others_ increases faster than *I*_self_ as *τ* grows (Fig. 4B,D), indicating that later PCs increasingly contain variance that makes individuals look more similar to one another. The difference is where individual information begins. At rest, identity is already present in the dominant component and is diluted by later components. During task, the leading component appears to be more group-shared, and identity emerges only after secondary directions are included. Thus, individual identity is not a fixed feature of a single FC representation. It is distributed across the manifold in a context-dependent way.

This has practical consequences in that if clinically or behaviorally relevant task differences reside in secondary PCs, scalar FC biomarkers may be insensitive to the relevant variance [7, 9]. Large samples are often required in brain-wide association studies because effect sizes are small [24]. One reason effect sizes may be small is that the representation itself attenuates behaviorally relevant dimensions. RV FC does not solve this problem automatically, but it gives a principled way to test whether the fingerprinting signal lives in dominant, intermediate, or high-variance explained PCA bands.

### 3.2 Hierarchical organization of within-parcel PCA structure

Within-parcel PC structure is hierarchically organized. As *τ* increases, each subspace nests the preceding one, and these progressively larger subspaces fall into a small number of ordered bands with distinct network and identifiability signatures, rather than varying smoothly. We interpret RV_*k*1_ − − RV_30_ as the dominant PCA band, RV_40_ − − RV_70_ as an secondary/intermediate band, and RV_80_ − − RV_90_ as a tertiary/high-variance residual band.

We use whitened SVD factors, giving each retained PC equal weight, so that RV measures the alignment between the subspaces spanned by two ROIs rather than being dominated by the few highest-variance directions [15]. This weighting allows later, lower-variance bands to contribute distinct information rather than being masked by the leading modes.

The evidence for these regimes is convergent rather than dependent on any single analysis. Between-threshold similarity matrices show block-like organization (Fig. 4A,C), while identifiability changes slope across similar thresholds (Fig. 4E). Network connectivity peaks at lower thresholds during rest and at higher thresholds during WM (Fig. 5). Parcel-identifiability rankings show self-similarity within regimes and, at rest, anticorrelation between the low- and high-threshold regimes (Fig. 6A(iv),B(iv)). The high-threshold regime exhibits low parcel identifiability in both conditions (Fig. 6A(iii),B(iii)), weak network structure at rest (Supplementary Fig. S2), and a limbic pattern consistent with physiological contamination. These regime boundaries should therefore be interpreted as approximate transition zones rather than fixed cutoffs.

Network organization reveals what the regimes contain. Rest RV_20_ shows high within-network and low between-network connectivity (Fig. 5A), consistent with canonical resting-state segregation [25]. WM RV_20_ shows both high within-network and high between-network connectivity (Fig. 5B), consistent with task-driven integration. At rest, network connectivity contracts at intermediate thresholds and expands again at high thresholds, especially between networks (Supplementary Fig. S3). This high-threshold expansion is driven by the limbic network, consistent with the vulnerability of nearby regions to cardiac and respiratory artifacts. At low thresholds, the visual network dominates within-network connectivity, reflecting coherent spontaneous activity in visual cortex. During WM, between-network structure persists deeper into the PC spectrum, even at RV_80_–RV_90_, whereas rest approaches a sparse disconnected graph (Supplementary Fig. S2). Task structure therefore appears to penetrate further into the regional manifold than resting-state structure.

### 3.3 Manifold-based connectivity

Strictly speaking, manifold-based connectivity represents a set of time series by a basis spanning a low-dimensional local subspace, with each edge measuring how strongly two subspaces align over time. In the PC case used here, RV can be interpreted as the normalized squared overlap between the temporal subspaces of two parcels. This turns FC from a scalar correlation between regional averages into a graph of subspace couplings. The resulting graph remains compatible with standard connectomic analysis, but its edges refer to relationships between local PCA representations rather than between single time series.

At the neuronal population scale, low-dimensional manifolds capture shared population dynamics and context-specific modes [1, 2]. At the whole-brain scale, PCA of parcel-averaged BOLD across tasks reveals a cognitive manifold in which leading and later components carry different task information [13]. Our results occupy an intermediate spatial scale: the PCA structure within each parcel, where the same organizing principle appears again. Dominant modes, secondary modes, and high-order residual modes are not interchangeable; they index different functional regimes. The band that carries individual identity can differ from the band that carries shared task structure.

Low variance-explained thresholds emphasize dominant dynamics. Intermediate thresholds expose secondary dynamics that may be invisible to scalar FC. High thresholds risk incorporating residual physiological, motion-related, or idiosyncratic variance. The appropriate threshold may therefore depend on the scientific question, as resting-state fingerprinting, task-related individual differences, group-level task effects, and residual-variance characterization may each occupy different portions of the PCA spectrum.

Although this study uses PCA because it provides orthonormal temporal bases and direct control over variance explained, the RV framework is more general. Any representation that yields comparable low-dimensional bases could be used to define subspace connectivity, including alternative linear decompositions or nonlinear embeddings after an appropriate basis representation.

RV could also be extended across spatial scales, for example to quantify coupling between parcel- and network-level representations, or across modalities with matched temporal structure. Neural activity is increasingly understood to exhibit multiscale organization [26], yet how latent representations at different scales relate to one another remains largely unexplored. Manifold-based connectivity provides a framework for asking how low-dimensional activity patterns align across levels of brain organization.

If brain activity is organized through nested low-dimensional subspaces, then connectomics should characterize how these subspaces align across regions, scales, cognitive states, and individuals. Our results show that this alignment varies systematically across the within-parcel PCA spectrum and follows recognizable spatial patterns of whole-brain organization. Across variance-explained thresholds, connectivity shifts along broad cortical hierarchies (Supplementary Figs. S6,S7); at rest, low-threshold structure is dominated by visual and somatomotor regions, whereas higher-threshold structure increasingly emphasizes association, default-mode, and limbic regions. A natural extension of this framework is to examine the eigenmodes of RV FC matrices, which may reveal additional low-dimensional principles of connectome organization [6].

### 3.4 Limitations

RV is strictly positive, so anti-correlations cannot be detected. However, Pearson R is overwhelmingly positive without global signal regression, and PCA already mean-centers within each parcel. No additional denoising was applied beyond HCP minimal preprocessing, so motion or physiological artifacts could differentially load onto different PCs. There is no ground truth for signal versus noise in the PC spectrum; the high-threshold interpretation rests on convergence across analyses and consistency with existing theory rather than direct artifact validation. Only the WM task was examined. Other tasks may shift regime boundaries, though we expect the general structure to hold. We do not make strong rest-versus-task comparisons given differences in scan duration, task design, and the use of phase-encoding direction as test–retest in WM.

## 4 Methods

### 4.1 Data

#### 4.1.1 Exploratory-Confirmatory Design

We first performed an exploratory analysis on the HCP Unrelated Subjects cohort (*n* = 100), then used the HCP S1200 New Subjects cohort for confirmatory analysis. After requiring complete, usable test–retest data, the confirmatory sample contained *n* = 191 subjects for resting state and *n* = 208 subjects for WM. We excluded subject sessions in which the first singular value captured nearly all within-ROI variance across every ROI (i.e., the data matrix was effectively rank-1 for all parcels), as such sessions are likely dominated by a global artifact such as head motion or respiratory contamination that would inflate between-subject similarity (*I*_others_) by imposing a shared covariance structure unrelated to neural activity. All analyses used previously collected, de-identified data from the Human Connectome Project. Ethics approval and informed consent for the original HCP data collection are described in Van Essen et al.[17]. No new participants were recruited and no new data were collected for the present study.

#### 4.1.2 HCP Task and Acquisition Details

Data were acquired on a customized 3T Siemens Connectome Skyra scanner with a 32channel head coil using a gradient-echo EPI sequence with multiband factor 8, TR = 720 ms, TE = 33.1 ms, flip angle = 52°, and 2.0 mm isotropic voxels (72 slices, FOV = 208 × 180 mm). Resting-state data consisted of four runs (REST1_LR, REST1_RL, REST2_LR, REST2_RL), each comprising 1200 timepoints (≈ 14.5 min), acquired with eyes open and fixation on a bright crosshair projected on a dark background. WM data consisted of two runs (WM_LR, WM_RL), each comprising 405 timepoints (≈ 4.9 min), using a block-design *N* -back task with 0-back and 2-back load conditions across four stimulus categories (faces, places, tools, body parts). Each run contained 8 task blocks (25 s each; 10 trials at 2.5 s per trial) interleaved with fixation blocks.

#### 4.1.3 Preprocessing

Data were minimally preprocessed using the HCP pipelines [18], which include gradient distortion correction, motion correction (6 DOF), fieldmap-based EPI distortion correction (TOPUP), boundary-based registration to T1w structural images, onestep spline resampling to MNI space, intensity normalization, and mapping of volume data to the fs_LR 32k cortical surface mesh via ribbon-constrained volume-to-surface projection, yielding CIFTI grayordinate time series (≈91,282 grayordinates: ≈59,412 cortical surface vertices and ≈31,870 subcortical voxels). No additional denoising (e.g., GSR, ICA-FIX, aCompCor) was applied. For resting state, each session’s FC was computed as the average across phase encoding directions (LR and RL); REST1 and REST2 served as test and retest sessions. For WM, LR and RL runs served as test and retest sessions.

#### 4.1.4 Parcellation

Cortical ROIs were defined using the Schaefer local-global parcellation [23] at the 100-, 200-, and 400-parcel resolutions, with parcels assigned to functional networks based on the 7-network parcellation from Yeo et al. [25].

### 4.2 RV Coefficient

The RV coefficient [27] is a multivariate generalization of Pearson correlation that can leverage principal component (PC) decompositions of voxel-level BOLD data within an ROI to construct FC matrices that progressively incorporate within-parcel spatial variation of BOLD while remaining computationally feasible. Specifically, we denote by RV_*τ*_ the RV-based FC matrix constructed from the PCs that collectively explain at least *τ* % of the within-ROI variance. RV quantifies the shared temporal dynamics between two ROIs’ principal component subspaces—the multivariate analog of how Pearson’s R quantifies shared dynamics between two scalar time series. In the univariate case, RV reduces to *R*^2^; thus, the framework contains the magnitude of Pearson coupling as its one-dimensional limit while extending it to local BOLD manifolds. The parcel average is itself a choice of representation; retaining multiple PCs is no less interpretable, and potentially more faithful to the underlying activity.

Early studies of RV FC by H. Zhang et al. used a seed-based FC “searchlight” approach, computing the RV coefficient between a fixed region of interest and a search cube [28, 29]. These approaches computed RV on voxel time series rather than PC time series. One could compute RV between all voxel time series in pairs of ROIs, but this creates the mathematical issue of correlating very large covariance subspaces, whose estimated association can be inflated by dimensionality independent of functional coupling. This motivates the use of low-dimensional PC subspaces. In the only other studies of the RV coefficient between ROI PCs for ROI-wise functional connectivity (“RV FC”), the variance explained threshold is taken as fixed [15, 16]. For instance, Ting et al. [15] retain PCs explaining up to 20% of the within-ROI variance and apply RV between the resulting low-dimensional representations to estimate resting-state network connectivity.

### 4.3 Overview

We begin with an overview of FC matrix construction. First, construct a voxel-by-time data matrix within an ROI and center each grayordinate time series across time before SVD. No variance standardization was applied before SVD. Then, use singular value decomposition to obtain principal component time series that are orthonormal in time, keeping the minimum number of principal components *q* whose sum of squared singular values over the sum of all squared singular values is ≥ *τ/*100. The RV coefficient is then computed between two ROIs’ PC time series as shown in Figure 1, with the procedure repeated for every edge. As with Pearson’s R, the absolute magnitude of RV is not directly interpretable; a value of 0.8 does not carry intrinsic meaning. Rather, RV values are informative in comparison—across thresholds, between subjects, or between conditions.

#### 4.3.1 Definition

Let **Y**(*t*) = (*Y*_*v*_(*t*))_*v*∈*S*_, where *Y*_*v*_(*t*) is the centered BOLD time series for voxel *v* and *S* is the set of all voxel indices. We partition the brain into *N* ROIs. For ROIs 1, …, *N*, let *S*_1_, …, *S*_*N*_ ⊂ *S* such that *S*_*i*_ ∩ *S*_*j*_ = ∅ for all *i* ≠ *j* and each *S*_*i*_ is the subset of voxel indices in ROI *i*. Collecting the centered voxel time series over *T* time points, the data matrix for ROI *i* is

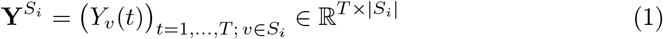

From singular value decomposition (SVD), we have

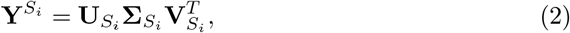

where 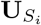 has orthonormal columns and represents temporal modes, 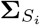 is the diagonal matrix of singular values, and 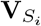 has orthonormal columns and represents spatial modes.

For the first *q*_*i*_ ≤ *r*_*i*_ singular components, where 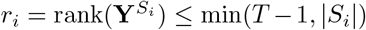 after temporal centering, we obtain a rank-*q*_*i*_ approximation of 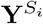. Using the singular values 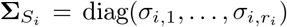, the minimum number of PCs *q*_*i*_ that explains at least *τ* % of the variance is chosen as 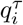 such that

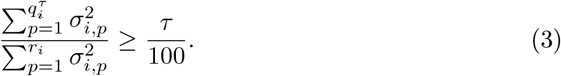

We denote the resulting RV-based FC matrix as RV_*τ*_ ; for example, RV_20_ retains PCs explaining at least 20% of within-ROI variance. Note that the two ROIs need not have the same number of PCs to reach a given *τ* .

In our implementation, we take the retained factor time series directly from the SVD, letting

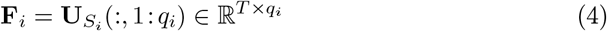

and defining **f**_*i*_(*t*) as the *t*-th row of **F**_*i*_. These are the “whitened” SVD factors. The factors are orthonormal in time, and **Q**_*i*_ collects the associated spatial patterns (absorbing the singular values). We use the local factor analysis (FA) model from Ting et al. [15]:

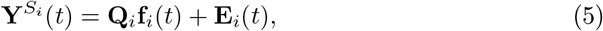

where 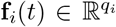 are latent factors summarizing the dynamics within ROI *i*, 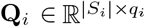 are the corresponding loadings, and **E**_*i*_(*t*) is residual noise.

To quantify the dependence between the latent structures of two ROIs *j* and *k*, we compute the RV coefficient between their factor series. Let 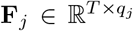 and 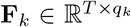 denote the matrices of retained factors for ROIs *j* and *k*. The standard RV coefficient is:

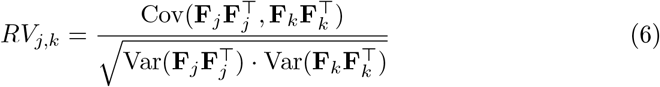

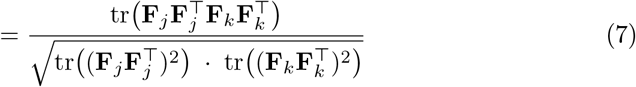

The equivalent and computationally efficient RV coefficient we use is:

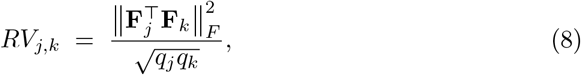

where ∥ · ∥_*F*_ denotes the Frobenius norm. We are able to obtain this simplification due to the orthonormality of the SVD-based factors. The denominator 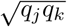 corrects for the dimensionality-dependent scaling of the numerator by accounting for the numbers of principal components retained in the two regional representations. Without this normalization, 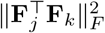 would tend to increase as more pairwise component alignments are included.

### 4.4 Identifiability

Identifiability, or the brain’s “fingerprint,” captures how well an individual’s functional connectome uniquely identifies them. Following Amico et al. [30], we define identifiability as follows. For a given *N*_ROI_ *× N*_ROI_ FC matrix FC_*i*_ for subject *i* among *N* subjects, assuming every subject has a test and retest session, let **A** ∈ R^*N ×N*^ be the identifiability matrix where *A*_*i,j*_ = Corr(FC_*i*_, FC_*j*_), with FC_*i*_ denoting the test-session FC matrix for subject *i* and FC_*j*_ the retest-session FC matrix for subject *j*. For the WM task, the test and retest sessions correspond to the two phase encoding directions (LR and RL), respectively. **A** is thus a non-symmetric square matrix. We define

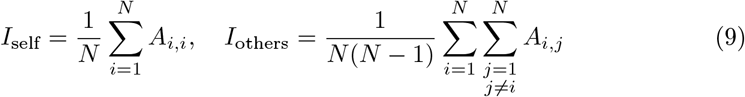

and

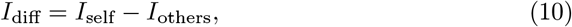

which quantifies how well an FC measure distinguishes individual brains from the population.

We consider parcel-level identifiability to understand the subject-specific information in the connectivity profile of ROI *r*. Let 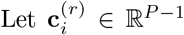 denote the vector of connections between ROI *r* and all other ROIs in subject *i*’s FC matrix (test session), and 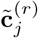 the analogous vector for the retest session of subject *j*. We define the parcel-level identifiability matrix **A**^(*r*)^ ∈ R^*N ×N*^ with entries 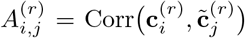.

Then

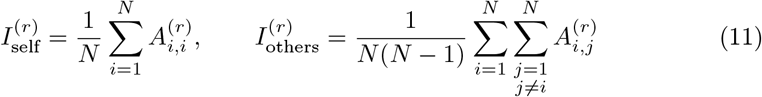

and

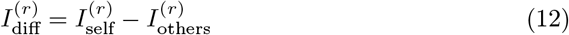

quantifies how well the connectivity profile of parcel *r* fingerprints individual subjects.

### 4.5 Statistical Analysis

#### 4.5.1 Replication and Robustness

Our findings replicated in the confirmatory sample. We quantified replication of the primary finding (Figure 4E) using the one-sided default Bayes factor for correlations (Rest: BF_10_ = 4.60 × 10^4^, *r* = 0.985; WM: BF_10_ = 1.64 × 10^3^, *r* = 0.953).

To assess robustness to parcellation granularity, we repeated the analysis using Schaefer 200 and 400 parcellations (Supplementary Figs. S20–S37). Within each condition, the relationship between variance explained threshold and identifiability was qualitatively preserved. At rest, the variance explained threshold *τ* at which RV-based identifiability drops below that of Pearson’s R shifted upward with finer parcellation, from approximately 20–30% at Schaefer 100 to 30–40% at Schaefer 200 and 40–50% at Schaefer 400. In WM, the inverted-U trajectory was likewise preserved, with identifiability peaking at intermediate variance thresholds rather than at either the scalar-like or high-variance extremes. Although the absolute number of principal components per parcel scales down with finer parcellations owing to fewer grayordinates per parcel, the same qualitative manifold decomposition is preserved across Schaefer 100, 200, and 400.

The correspondence between R and RV matrices, quantified by *r*(*R*, RV_*τ*_ ), declined monotonically with increasing *τ* for rest across all three parcellations (e.g., at *k*_1_: *r* = 0.92, 0.92, 0.90 for Schaefer 100, 200, 400, respectively), with BF_10_ *>* 100 at every threshold and parcellation tested. Identifiability at the lowest thresholds (*k*_1_, 10%) drifted upward with finer parcellations (peak rest *I*_diff_: ≈ 0.22, 0.23, 0.26 for Schaefer 100, 200, 400), likely reflecting the increased spatial specificity of smaller parcels. Despite this upward drift at low thresholds, the monotonic decline in rest *I*_diff_ with increasing *τ* was preserved across all three parcellations, as was the invertedU trajectory in WM. The crossover threshold at which RV-based *I*_diff_ falls below Pearson *I*_diff_ in rest shifted upward with finer parcellations (*τ* ≈ 20–30% at Schaefer 100; *τ* ≈ 30–40% at Schaefer 200; *τ* ≈ 40–50% at Schaefer 400), consistent with the fact that finer parcellations have fewer grayordinates per parcel and thus require a higher *τ* to accumulate the same number of PCs. That is, despite the shift in *τ*, the number of principal components per parcel at the crossover is comparable across parcellations, indicating that the transition from RV-dominant to R-dominant identifiability occurs at a consistent level of within-parcel dimensionality regardless of parcellation resolution. The *r*(*R*, RV_*τ*_ ) values were also systematically higher at finer parcellations for matched *τ* (e.g., at RV_80_: *r* = 0.14, 0.32, 0.45 for Schaefer 100, 200, 400), indicating that finer parcellations retain more R-like structure at higher variance thresholds.

#### 4.5.2 One-Sided Default Bayes Factor for Correlations

We quantify evidence for correlation using a one-sided default Bayes factor for the correlation coefficient, following Ly et al. [31]. We test *H*_1_: *r >* 0 versus *H*_0_: *r* = 0 (no correlation). The Bayes factor BF_10_ is the ratio of marginal likelihoods under *H*_1_ and *H*_0_:

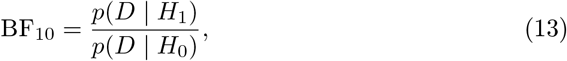

where *D* denotes the observed data (the paired exploratory and confirmatory values). Under *H*_1_, we placed a default prior on *r* supported on [0, 1] (one-sided). The computation uses the standard Jeffreys prior for the correlation coefficient and the hypergeometric function _2_*F*_1_ for the marginal likelihood; refer to Ly et al. [31] for more details. BF_10_ *>* 1 indicates evidence for replication (positive correlation); BF_10_ *<* 1 indicates evidence against replication.

#### 4.5.3 Bayes Factor Interpretation

We report Bayes factors in favor of the alternative hypothesis over the null, denoted BF_10_. Thus BF_10_ *>* 1 indicates evidence for the alternative, and BF_10_ *<* 1 indicates evidence for the null (with BF_01_ = 1*/*BF_10_). We interpreted BF_10_ using the scale proposed by Lee and Wagenmakers [32]:

- 1 *<* BF_10_ ≤ 3: anecdotal evidence for *H*_1_
- 3 *<* BF_10_ ≤ 10: moderate evidence for *H*_1_
- 10 *<* BF_10_ ≤ 30: strong evidence for *H*_1_
- 30 *<* BF_10_ ≤ 100: very strong evidence for *H*_1_
- BF_10_ *>* 100: decisive evidence for *H*_1_

The inverse scale applies when BF_10_ *<* 1 (e.g., 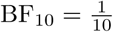 provides moderate evidence for *H*_0_).

## Data availability

The resting-state and working-memory fMRI data used in this study were obtained from the Human Connectome Project Young Adult dataset, including the HCP 100 Unrelated Subjects and S1200 releases. These data are available through ConnectomeDB after registration and acceptance of the WU-Minn Human Connectome Project Data Use Terms. Aggregate numerical source data underlying the figures are available at https://doi.org/10.5281/zenodo.21343846. These released source-data tables contain only aggregate derived values and do not include raw HCP data, HCP subject identifiers, exploratory or confirmatory subject lists, subject-level feature matrices, or restricted-access variables. Reuse of the released aggregate tables should follow the applicable WU-Minn HCP Open Access Data Use Terms.

## Code availability

Code used to construct the RV-based functional connectivity matrices, calculate connectome identifiability, and reproduce the analyses and figures reported in this study is available at https://github.com/nathoogroup/fingerprintrv. An archived version of the code used for this manuscript is available at https://doi.org/10.5281/zenodo.21343846.

## Acknowledgements

Data were provided by the Human Connectome Project, WU–Minn Consortium (Principal Investigators: David Van Essen and Kamil Ugurbil; 1U54MH091657), funded by the 16 NIH Institutes and Centers that support the NIH Blueprint for Neuroscience Research and by the McDonnell Center for Systems Neuroscience at Washington University. ChatGPT (OpenAI) was used to assist in identifying potentially relevant literature and copyediting (formatting, proof-reading, and minor wording suggestions for clarity). All retrieved references are independently verified by the authors, and all screening, interpretation, and citation decisions are made by the authors.

## Author contributions

Conceptualization: P.V., E.A., F.S.N., and M.F.M.; Methodology: P.V., E.A., F.S.N., and M.F.M.; Software: P.V.; Validation: P.V.; Formal analysis: P.V.; Investigation: P.V.; Data curation: P.V.; Visualization: P.V.; Writing–original draft: P.V.; Writing– review and editing: P.V., E.A., F.S.N., and M.F.M.; Supervision: E.A., F.S.N., and M.F.M.; Project administration: F.S.N. and M.F.M. All authors reviewed and approved the submitted manuscript.

