## Supplemental Information for "Neural manifold connectomics reveals multiregime functional connectivity"

### Supplementary Information

#### Supplementary Figures

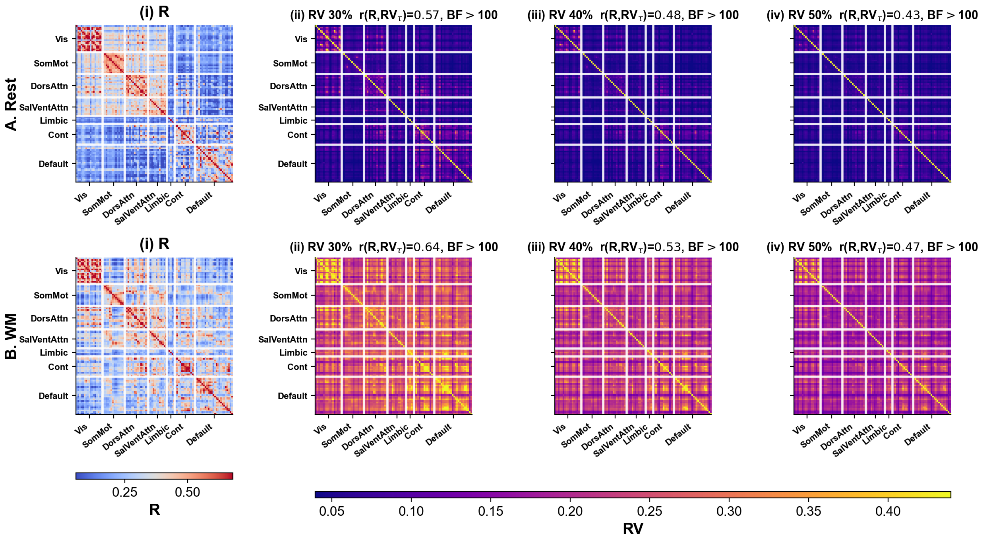


**Supplementary Fig. S1**: **Group FC matrices at moderate variance thresholds (Schaefer 100, confirmatory sample).** A. Rest; B. WM. i. Pearson's R-based FC. RV-based FC matrices retaining PCs that explain at least ii. 30%, iii. 40%, iv. 50% of the within-ROI variance. Matrices are network-reordered by the Yeo 7-network parcellation. $r(R, \mathrm{RV}_\tau)$ and one-sided default Bayes factors ($\mathrm{BF}_{10}$, $H_1$: $r > 0$) are shown for each threshold. All $\mathrm{BF}_{10} > 100$. Extends main text Figure 1.

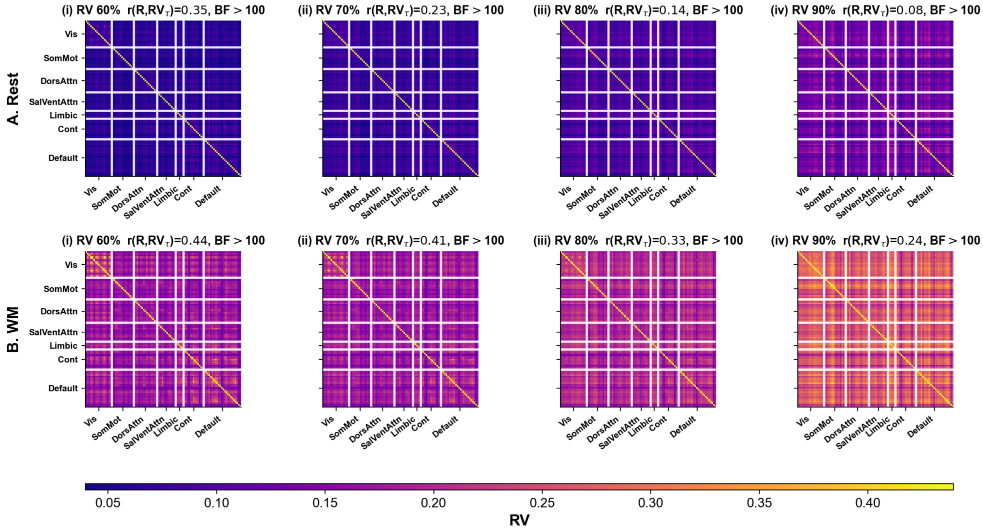


**Supplementary Fig. S2**: **Group FC matrices at high variance thresholds (Schaefer 100, confirmatory sample).** A. Rest; B. WM. RV-based FC matrices retaining PCs that explain at least i. 60%, ii. 70%, iii. 80%, iv. 90% of the within-ROI variance. Network block-diagonal structure becomes increasingly prominent, particularly in WM, while rest matrices become increasingly sparse. All $BF_{10} > 100$. Extends main text Figure 1.

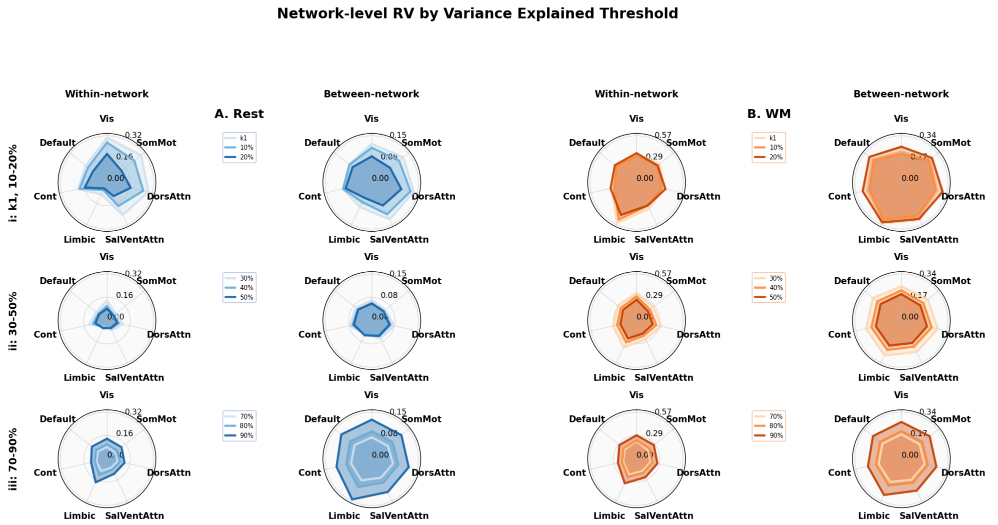


**Supplementary Fig. S3**: **Network-level RV across all variance thresholds (Schaefer 100, confirmatory sample).** Yeo 7 network radar plots for group RV-based FC across variance bands. A. Rest; B. WM. i. Within-network mean RV; ii. Between-network mean RV. Rows: i. $k_1$, 10–20%; ii. 30–50%; iii. 70–90%. Contours darken with increasing variance explained threshold. Within-network connectivity contracts from low to moderate thresholds before expanding again at high thresholds, driven primarily by limbic network activation. Extends main text Figure 4.

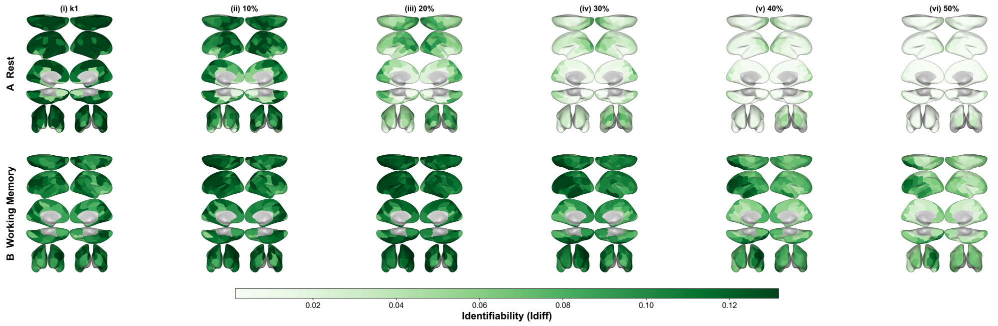


**Supplementary Fig. S4**: **Parcel-level identifiability across low-to-moderate thresholds (Schaefer 100, confirmatory sample).** A. Rest; B. WM. Columns show $I_{\text{diff}}^{(r)}$ from i. $\text{RV}_{k1}$, ii. $\text{RV}_{10}$, iii. $\text{RV}_{20}$, iv. $\text{RV}_{30}$, v. $\text{RV}_{40}$, vi. $\text{RV}_{50}$ FC. Color scale: parcel-level differential identifiability, darker green = higher. Identifiability is high and broadly distributed at low thresholds, becoming increasingly sparse at moderate thresholds. Extends main text Figure 5.

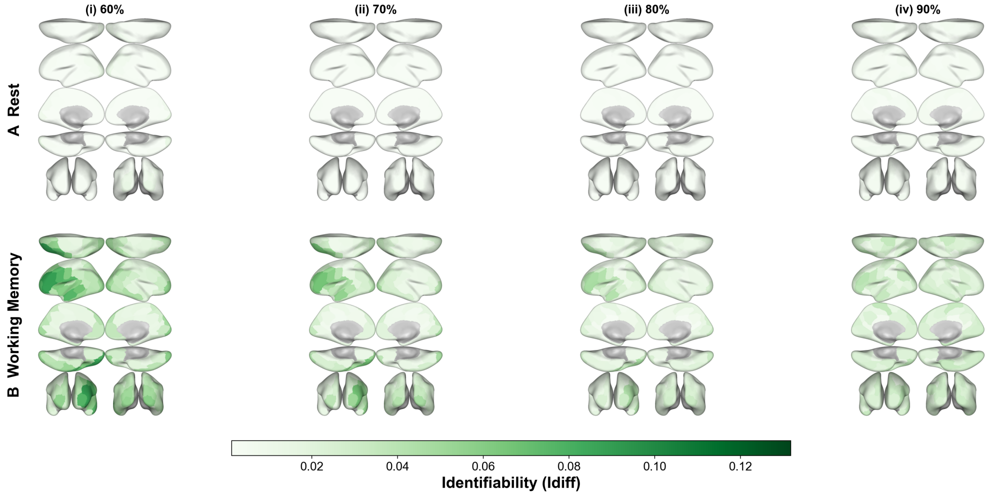


**Supplementary Fig. S5**: **Parcel-level identifiability across high thresholds (Schaefer 100, confirmatory sample).** A. Rest; B. WM. Columns show $I^{(r)}_{\text{diff}}$ from i. $\text{RV}_{60}$, ii. $\text{RV}_{70}$, iii. $\text{RV}_{80}$, iv. $\text{RV}_{90}$ FC. At high thresholds, identifiability is uniformly low in rest; residual identifiability in WM concentrates in limbic and default mode regions. Extends main text Figure 5.

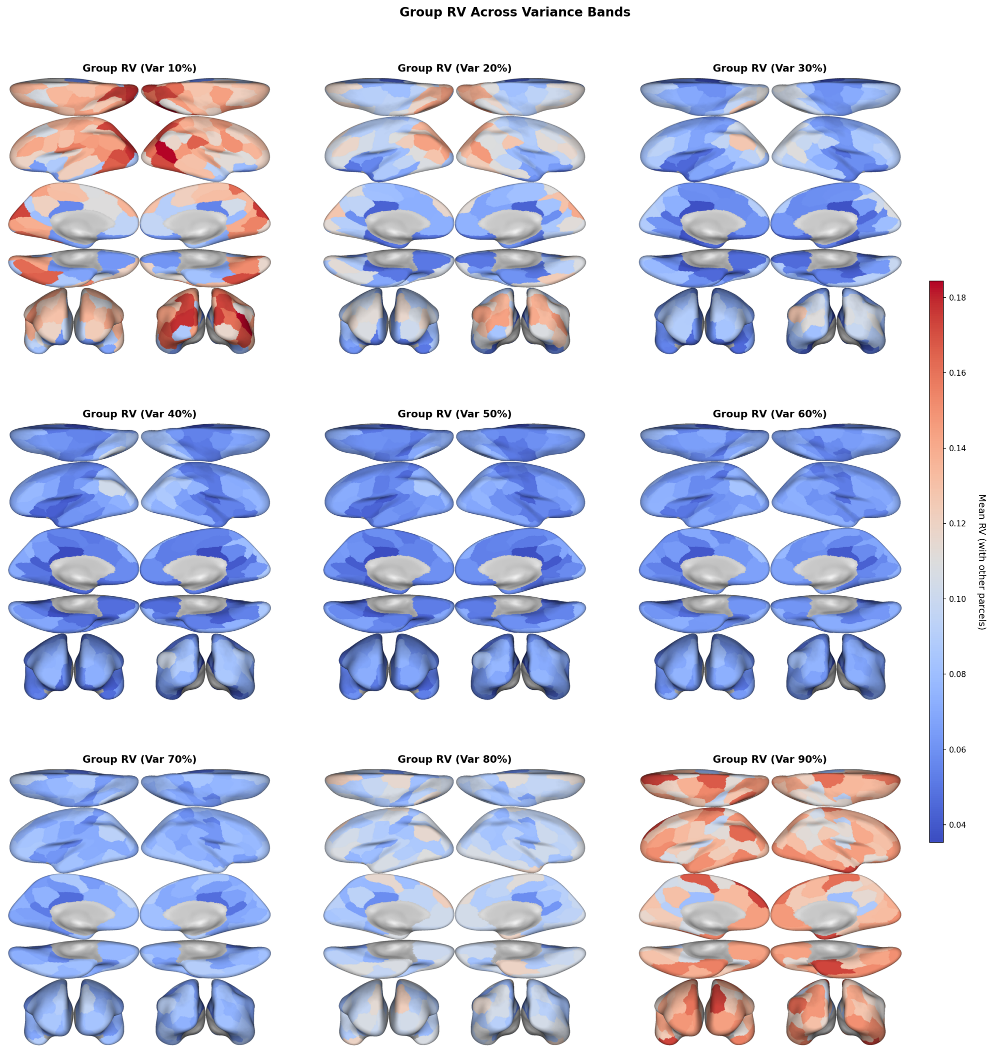


**Supplementary Fig. S6**: **Parcel-level group RV brain maps across cumulative variance thresholds (Rest, Schaefer 100, confirmatory sample).** Each panel shows the row-averaged group RV matrix at a cumulative variance threshold, computed as the mean RV between each parcel and all other parcels. Thus, $RV_{20}$ includes the PCs needed to explain at least 20% of within-ROI variance, $RV_{60}$ includes PCs through 60%, and so on, rather than isolating adjacent variance bands.

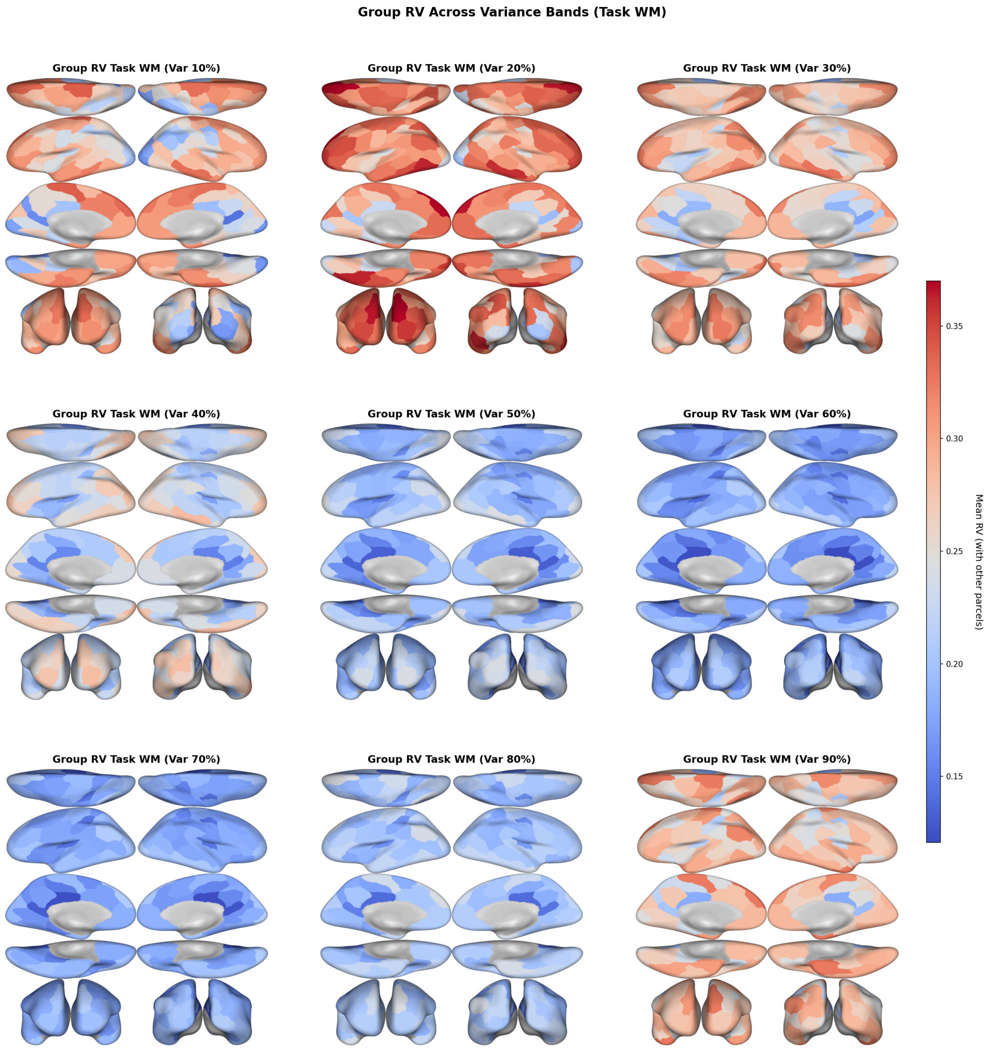


**Supplementary Fig. S7**: **Parcel-level group RV brain maps across cumulative variance thresholds (WM, Schaefer 100, confirmatory sample).** Same layout as the resting-state cumulative RV brain maps for the working memory task. Parcel values are row means of the cumulative group RV matrix, averaging each parcel's RV with all other parcels at each variance threshold.

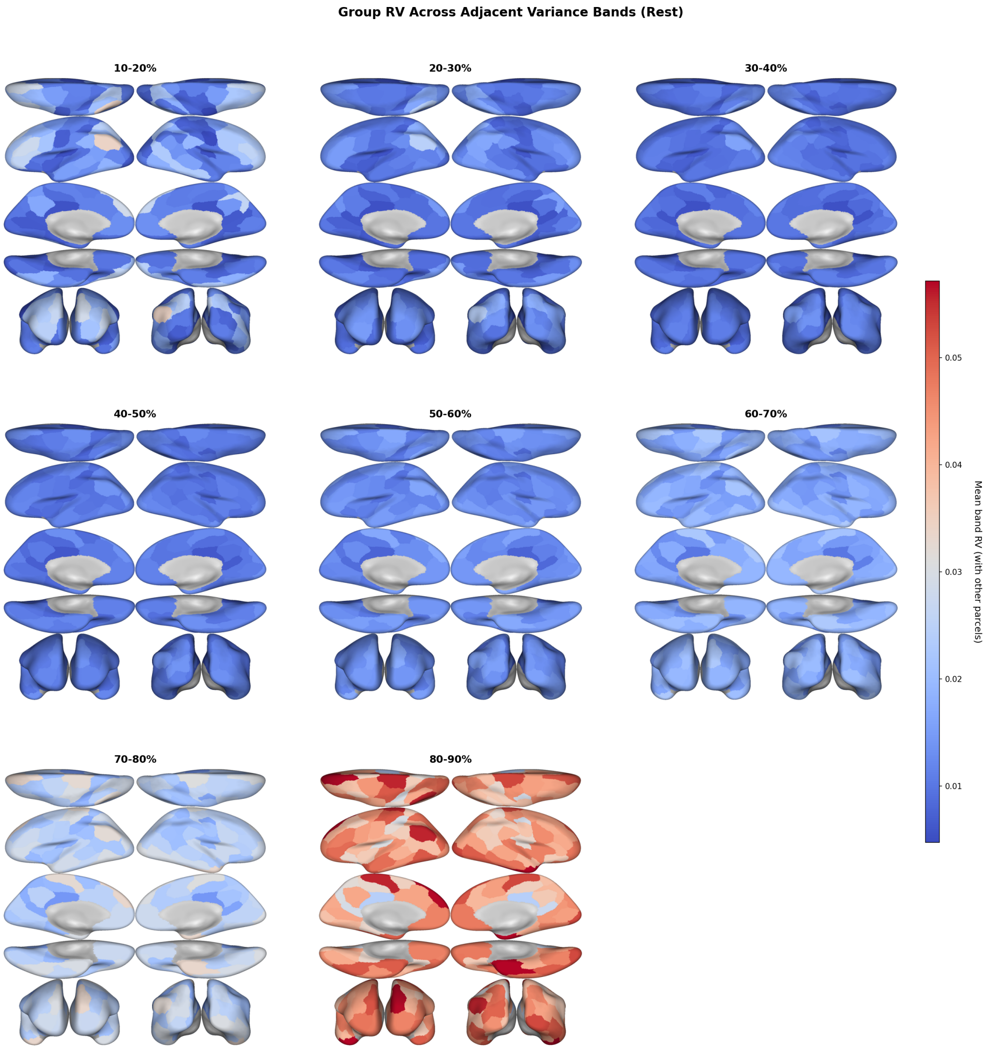


**Supplementary Fig. S8**: **Group RV brain maps across adjacent variance bands (Rest, Schaefer 100, confirmatory sample).** Each panel shows the mean RV of each parcel (averaged across all edges) for PCs explaining variance within a specific band: 10–20%, 20–30%, . . . , 80–90%. Early bands show high RV concentrated in visual and somatomotor cortex; later bands show a shift toward higher RV in frontoparietal, default mode, and limbic regions.

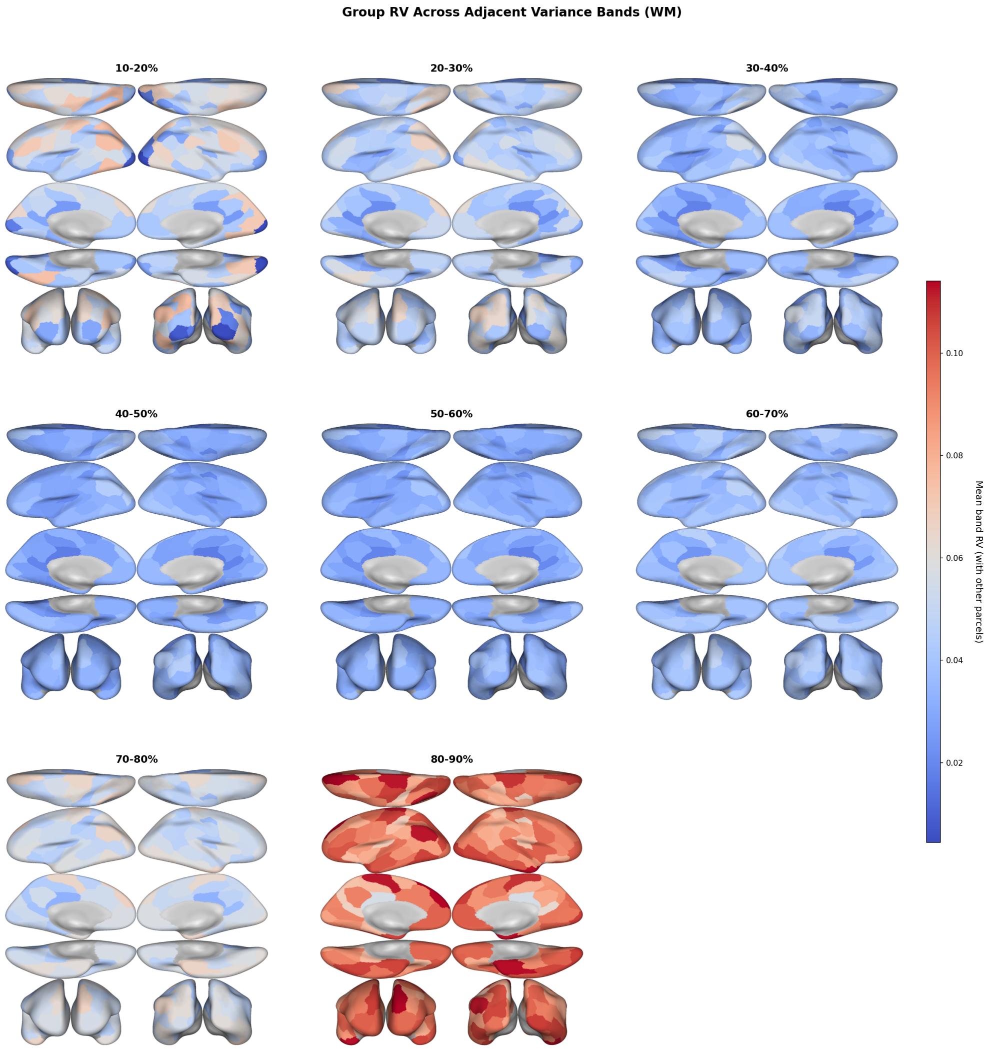


**Supplementary Fig. S9**: **Group RV brain maps across adjacent variance bands (WM, Schaefer 100, confirmatory sample).** Same layout as the resting-state adjacent-band brain maps for the working memory task.

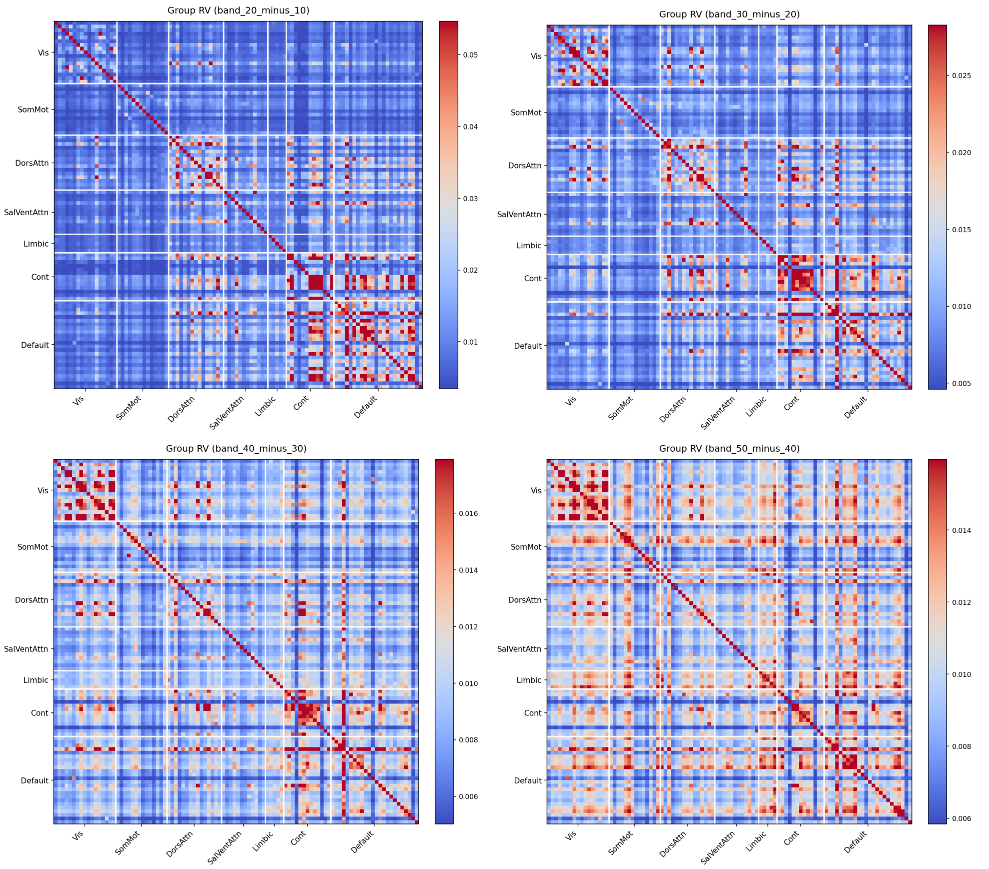


**Supplementary Fig. S10**: **Group band RV matrices, low-to-moderate bands (Rest, confirmatory sample).** Network-reordered group-averaged RV matrices computed from PCs within adjacent variance bands. Top left: 10–20%; top right: 20–30%; bottom left: 30–40%; bottom right: 40–50%. These FC matrices are constructed from the PCs that explain the *additional* variance between consecutive thresholds, isolating the contribution of each band to the overall connectivity structure.

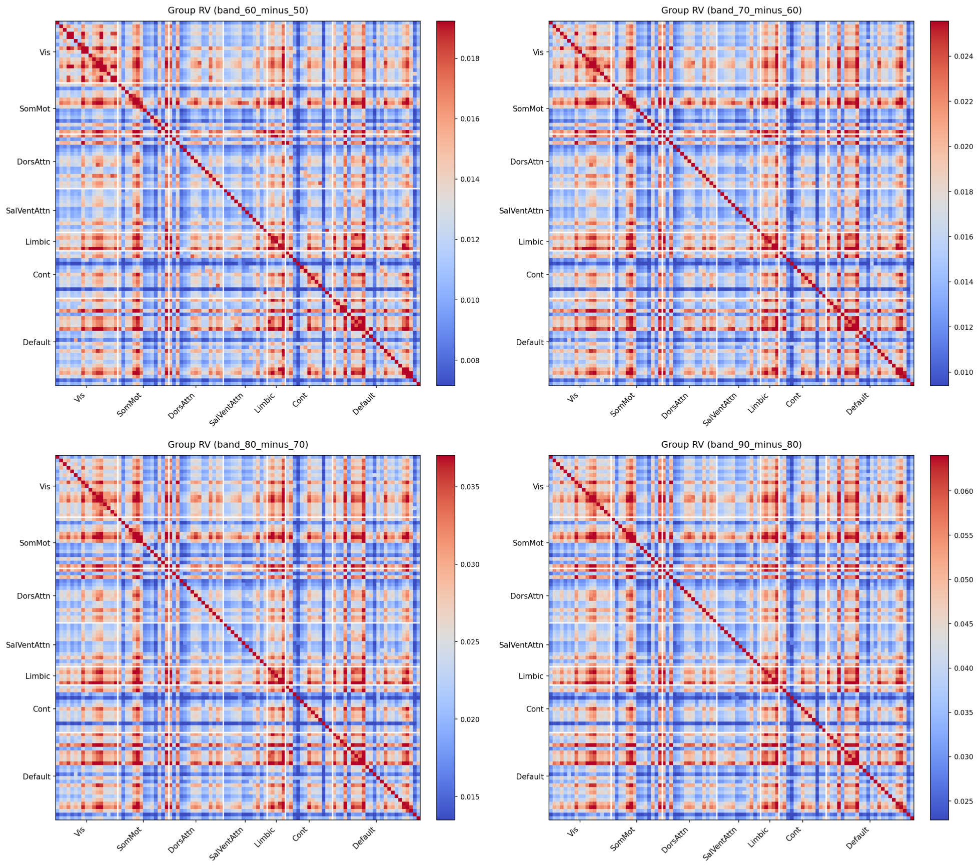


**Supplementary Fig. S11**: **Group band RV matrices, moderate-to-high bands (Rest, confirmatory sample).** Top left: 50–60%; top right: 60–70%; bottom left: 70–80%; bottom right: 80–90%. At higher bands, connectivity becomes increasingly sparse with residual structure concentrated along the block diagonal, consistent with network-specific fine-grained voxel-level variation.

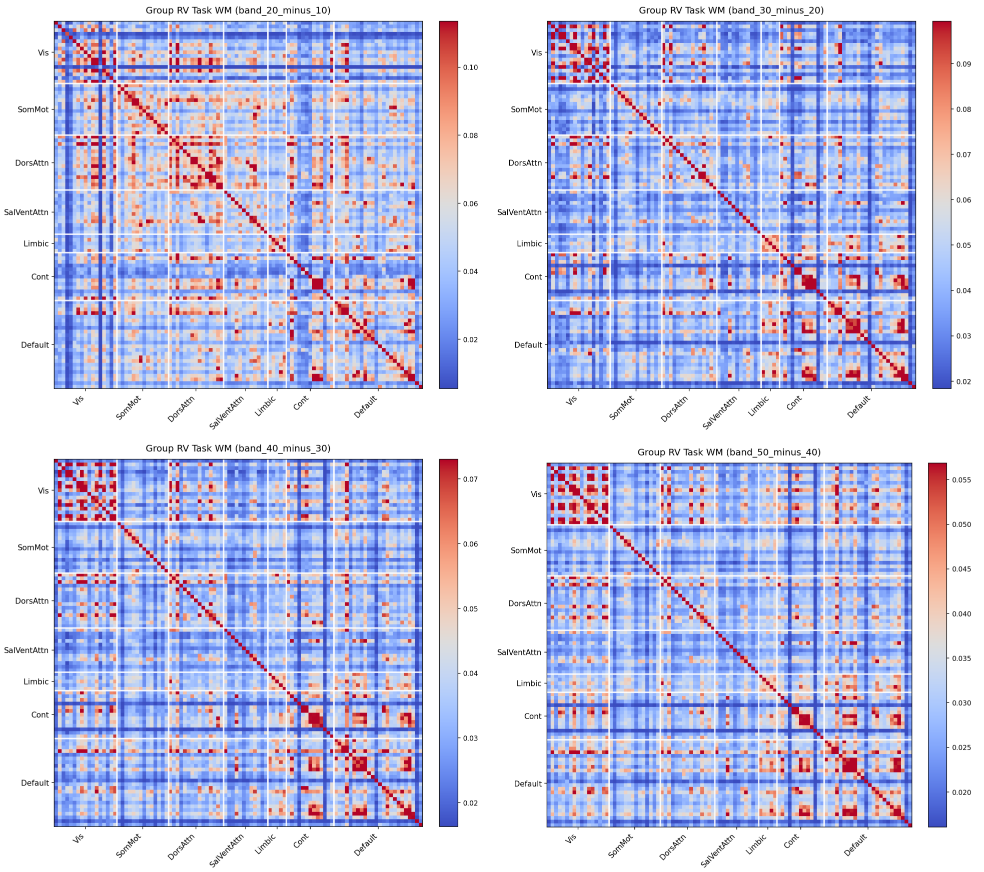


**Supplementary Fig. S12: Group band RV matrices, low-to-moderate bands (WM, confirmatory sample).** Same layout as the resting-state low-to-moderate band matrices for the working memory task. Top left: 10–20%; top right: 20–30%; bottom left: 30–40%; bottom right: 40–50%.

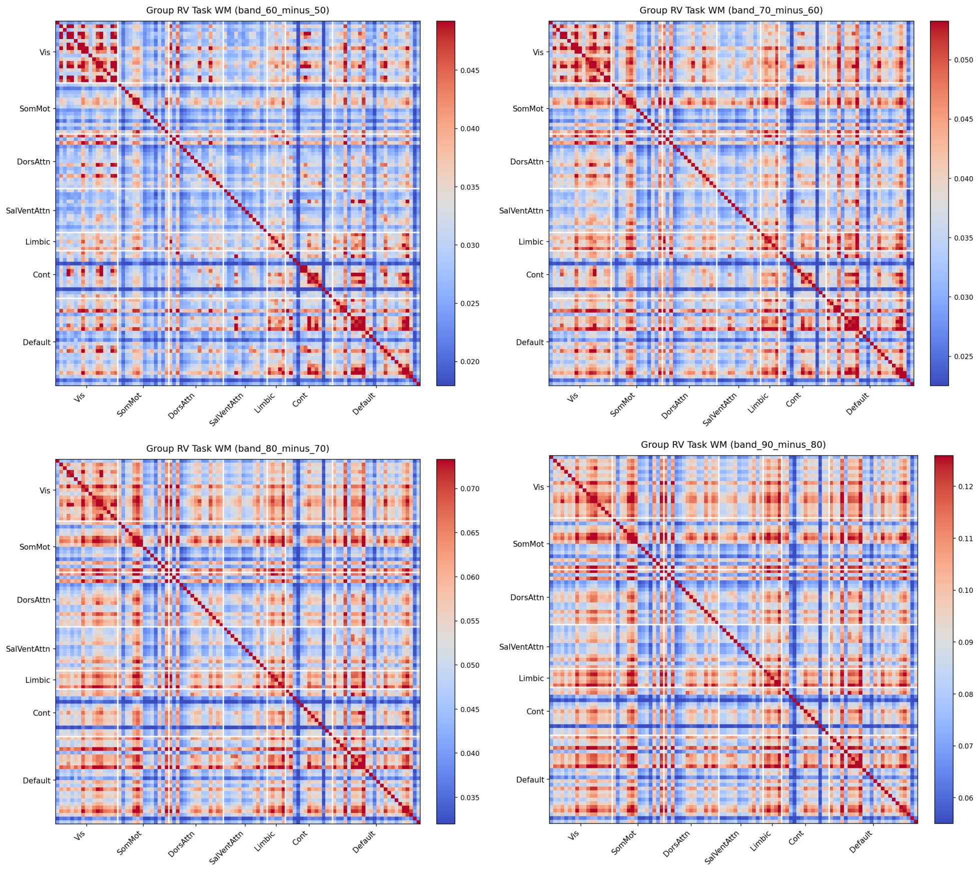


**Supplementary Fig. S13**: **Group band RV matrices, moderate-to-high bands (WM, confirmatory sample).** Same layout as the resting-state moderate-to-high band matrices for the working memory task. Top left: 50–60%; top right: 60–70%; bottom left: 70–80%; bottom right: 80–90%.

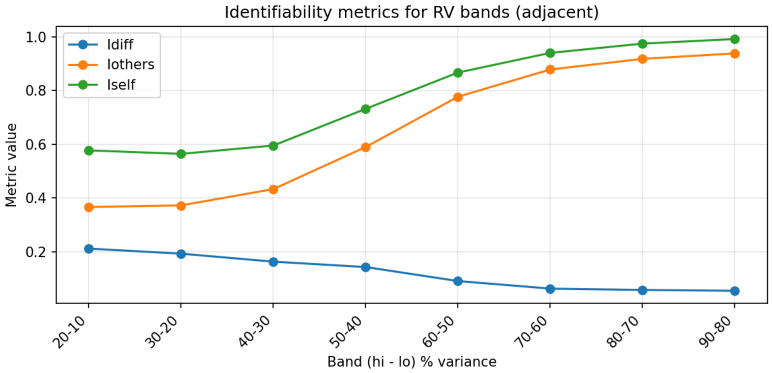


**Supplementary Fig. S14**: **Identifiability metrics across adjacent variance bands (Rest, confirmatory sample).** $I_{\mathrm{diff}}$, $I_{\mathrm{others}}$, and $I_{\mathrm{self}}$ as a function of adjacent variance band (10–20% through 80–90%). $I_{\mathrm{self}}$ and $I_{\mathrm{others}}$ increase monotonically with band, while $I_{\mathrm{diff}}$ remains relatively stable before slightly decreasing at the highest bands, indicating that later PCs contribute increasingly shared (non-individual) variance.

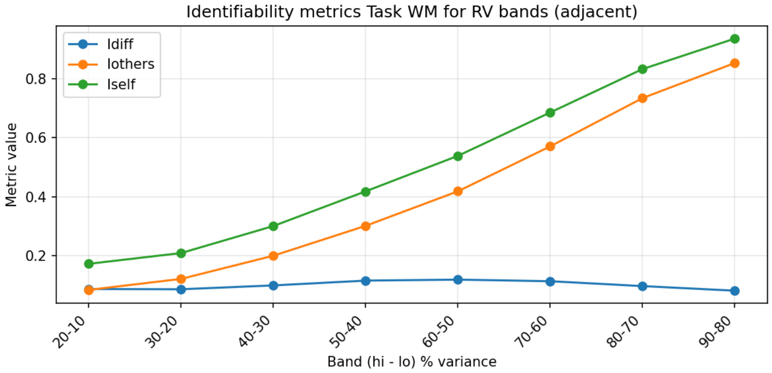


**Supplementary Fig. S15**: **Identifiability metrics across adjacent variance bands (WM, confirmatory sample).** Same layout as the resting-state adjacent-band identifiability metrics for the working memory task.

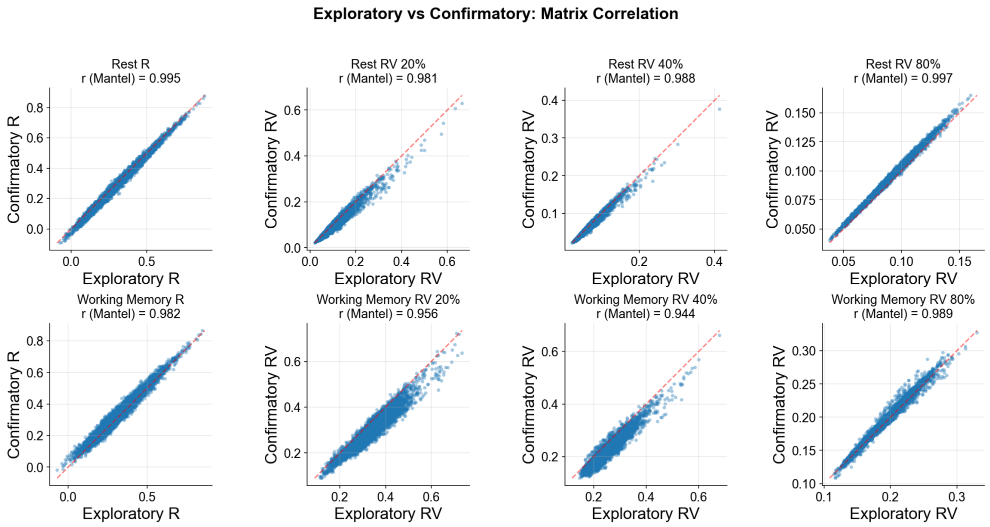


**Supplementary Fig. S16**: **Exploratory vs. confirmatory group FC matrix correlation.** Scatter plots of upper-triangular entries of exploratory group FC matrices ($x$-axis) against confirmatory group FC matrices ($y$-axis) for Pearson's R and $\text{RV}_{20}$, $\text{RV}_{40}$, $\text{RV}_{80}$ FC. Top: Rest; bottom: WM. Mantel $r$ values shown for each panel. All correlations are strongly positive (Rest R: $r = 0.995$; Rest $\text{RV}_{20}$: $r = 0.981$; Rest $\text{RV}_{40}$: $r = 0.988$; Rest $\text{RV}_{80}$: $r = 0.997$; WM R: $r = 0.982$; WM $\text{RV}_{20}$: $r = 0.956$; WM $\text{RV}_{40}$: $r = 0.944$; WM $\text{RV}_{80}$: $r = 0.989$), indicating that group-level FC structure is highly reproducible across independent samples at all thresholds.

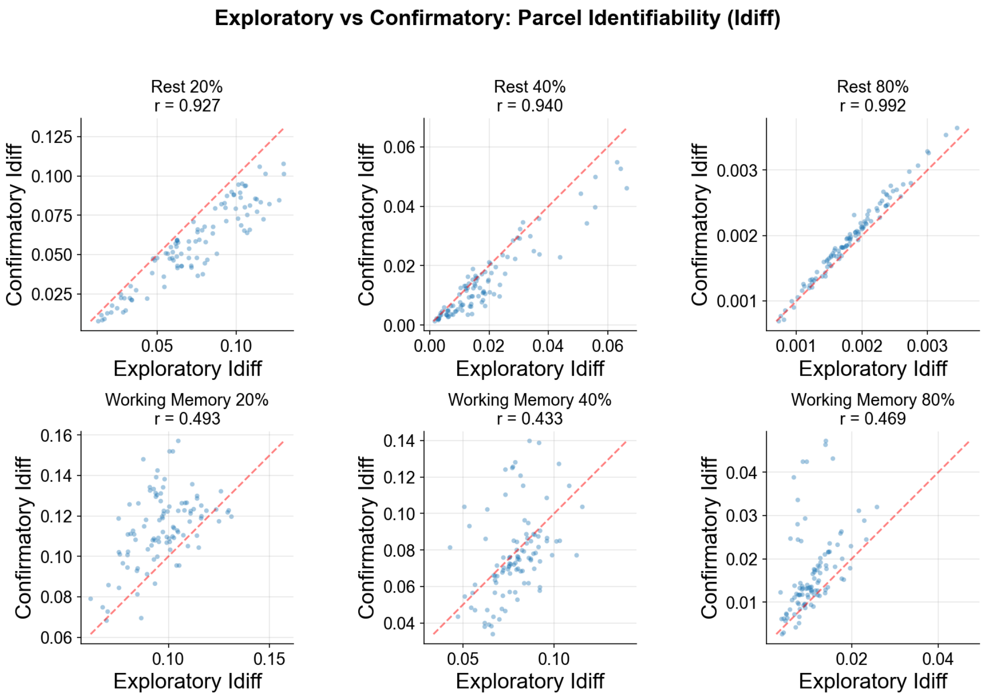


**Supplementary Fig. S17**: **Exploratory vs. confirmatory parcel-level $I^{(r)}_{\text{diff}}$ correlation.** Scatter plots of parcel-level $I^{(r)}_{\text{diff}}$ in the exploratory sample ($x$-axis) against the confirmatory sample ($y$-axis) at $\text{RV}_{20}$, $\text{RV}_{40}$, $\text{RV}_{80}$. Top: Rest ($r$ = 0.927, 0.940, 0.992); bottom: WM ($r$ = 0.493, 0.433, 0.469). Resting-state parcel identifiability replicates strongly; WM replication is moderate, consistent with the noisier parcel-level identifiability in task conditions.

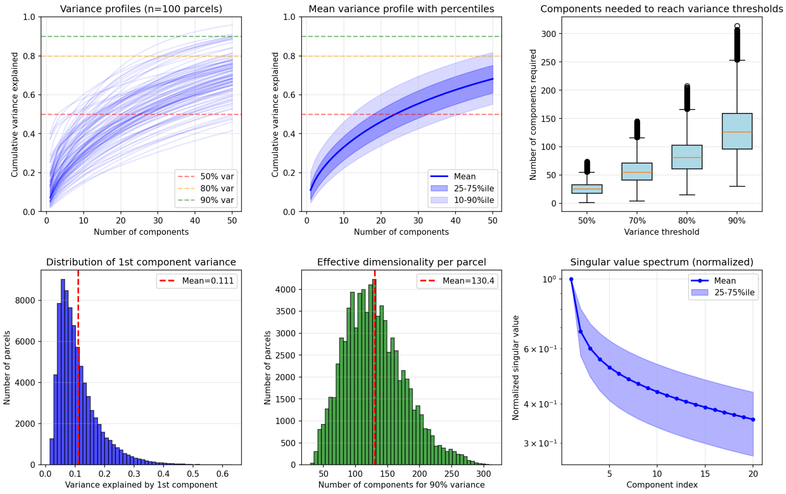


**Supplementary Fig. S18**: **Variance explained profiles (Rest, confirmatory sample).** Distribution of cumulative variance explained by the first $q$ PCs across parcels and subjects in resting-state data.

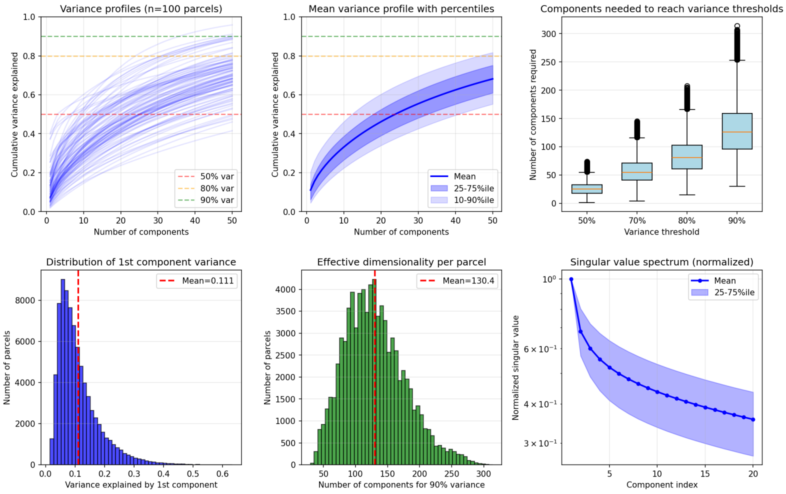


**Supplementary Fig. S19**: **Variance explained profiles (WM, confirmatory sample).** Distribution of cumulative variance explained by the first $q$ PCs across parcels and subjects in working memory data.

#### Schaefer 200 parcellation

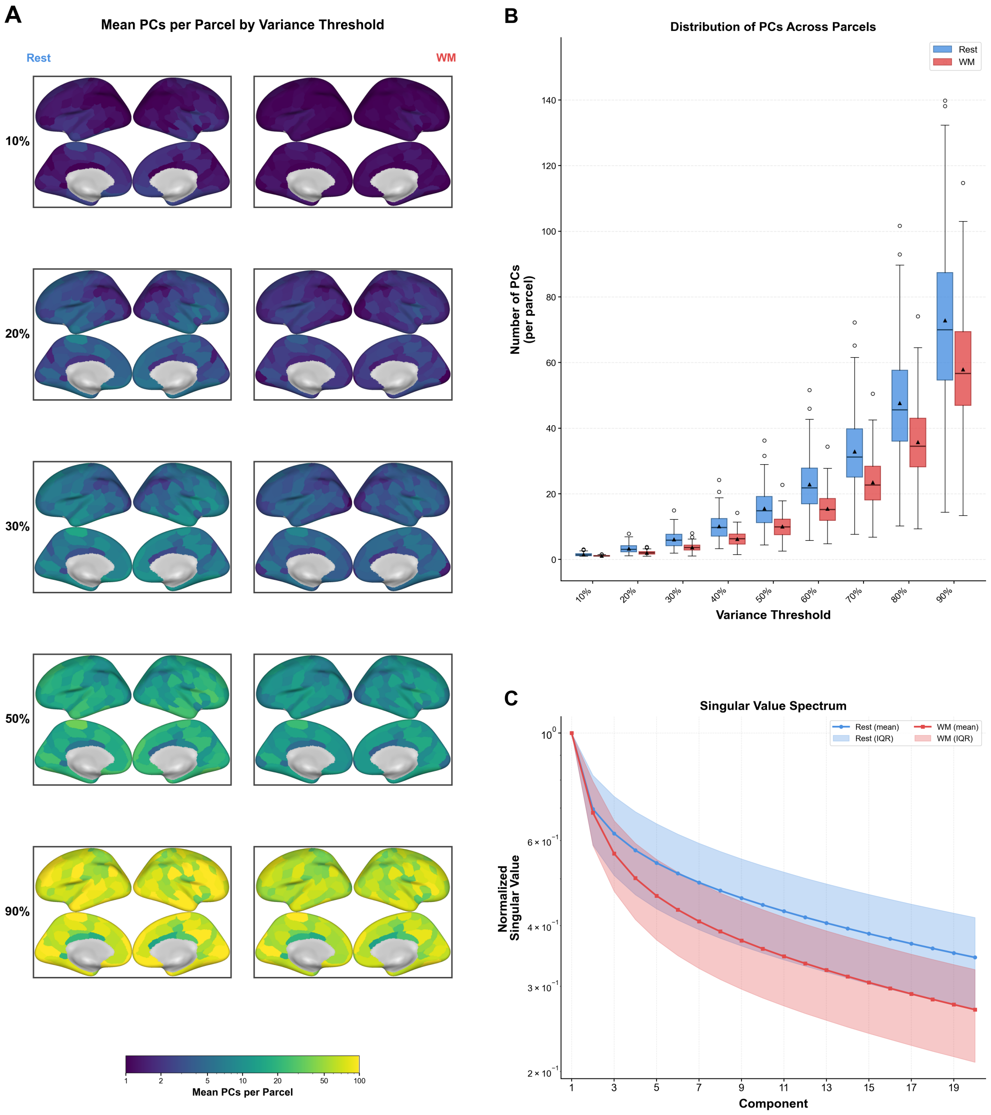


**Supplementary Fig. S20**: **PCs needed to explain within-ROI variance (Schaefer 200, confirmatory sample).** Same layout as main text Figure 2 at the 200-parcel resolution. A. Mean PCs per parcel by variance threshold for Rest (left) and WM (right). B. Distribution of PCs across parcels at each threshold. C. Singular value spectrum (mean and IQR). The spatial heterogeneity of within-ROI dimensionality is preserved at finer parcellation, with occipito-parietal ROIs requiring more PCs than frontal ROIs. Absolute PC counts are lower than Schaefer 100 owing to fewer grayordinates per parcel.

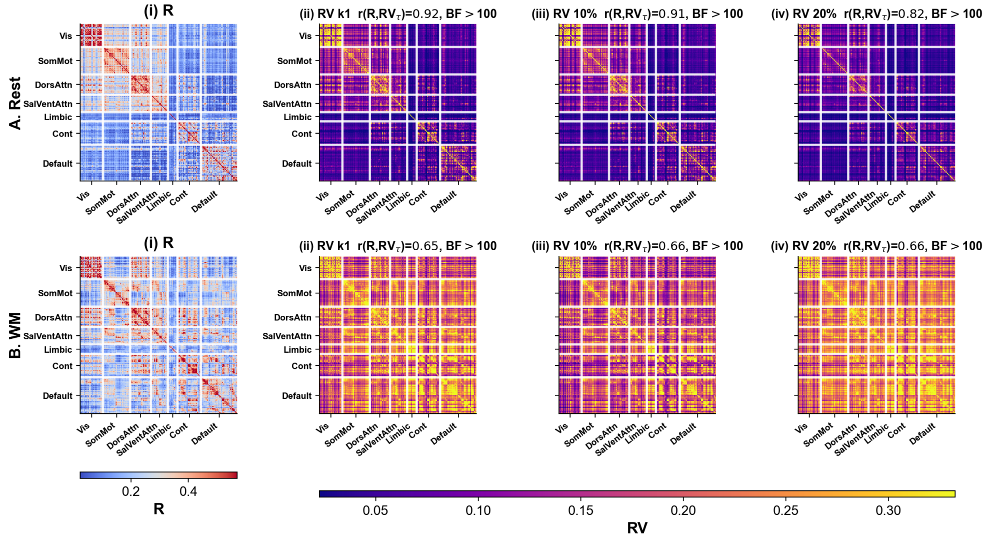


**Supplementary Fig. S21**: **Group FC matrices at low variance thresholds (Schaefer 200, confirmatory sample).** A. Rest; B. WM. i. Pearson's R. RV-based FC at ii. $k_1$, iii. 10%, iv. 20% variance explained. $r(R, \mathrm{RV}_\tau)$ and $\mathrm{BF}_{10}$ shown for each threshold. The high correspondence between R and $\mathrm{RV}_{k1}$ ($r = 0.92$) replicates the Schaefer 100 finding.

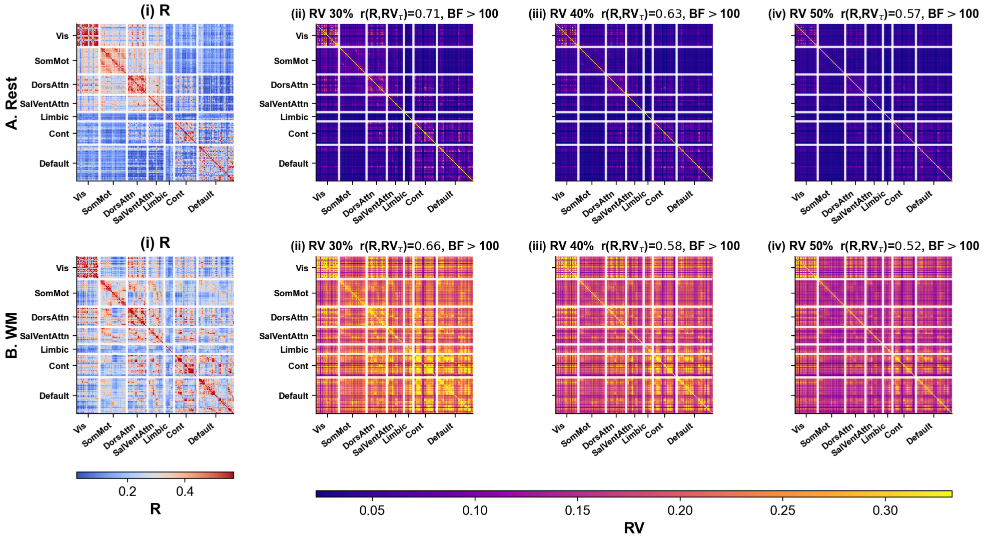


**Supplementary Fig. S22**: **Group FC matrices at moderate variance thresholds (Schaefer 200, confirmatory sample).** A. Rest; B. WM. i. Pearson's R. RV-based FC at ii. 30%, iii. 40%, iv. 50% variance explained. $r(R, \mathrm{RV}_\tau)$ values are systematically higher than at the corresponding thresholds for Schaefer 100, indicating that finer parcellations retain more R-like structure.

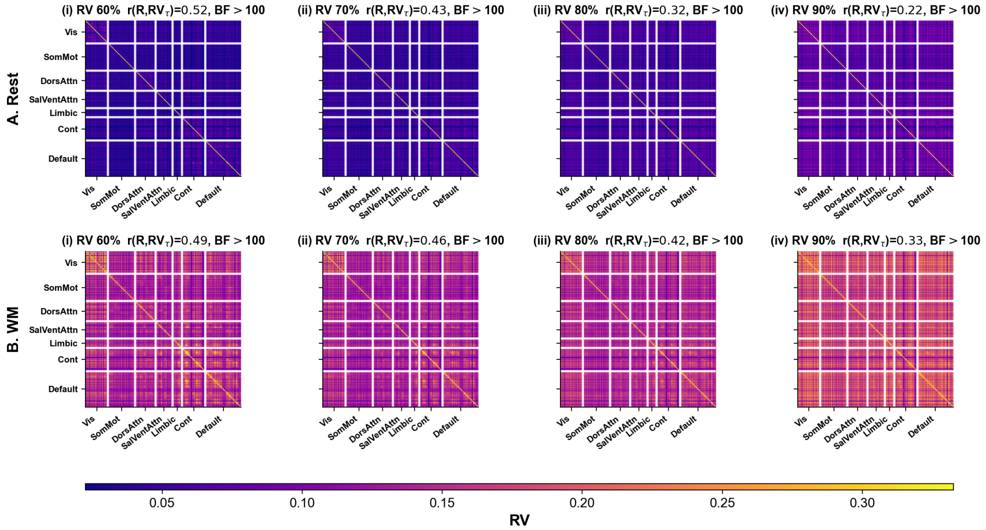


**Supplementary Fig. S23**: **Group FC matrices at high variance thresholds (Schaefer 200, confirmatory sample).** A. Rest; B. WM. RV-based FC at i. 60%, ii. 70%, iii. 80%, iv. 90% variance explained. Network block-diagonal structure emerges at comparable thresholds as Schaefer 100.

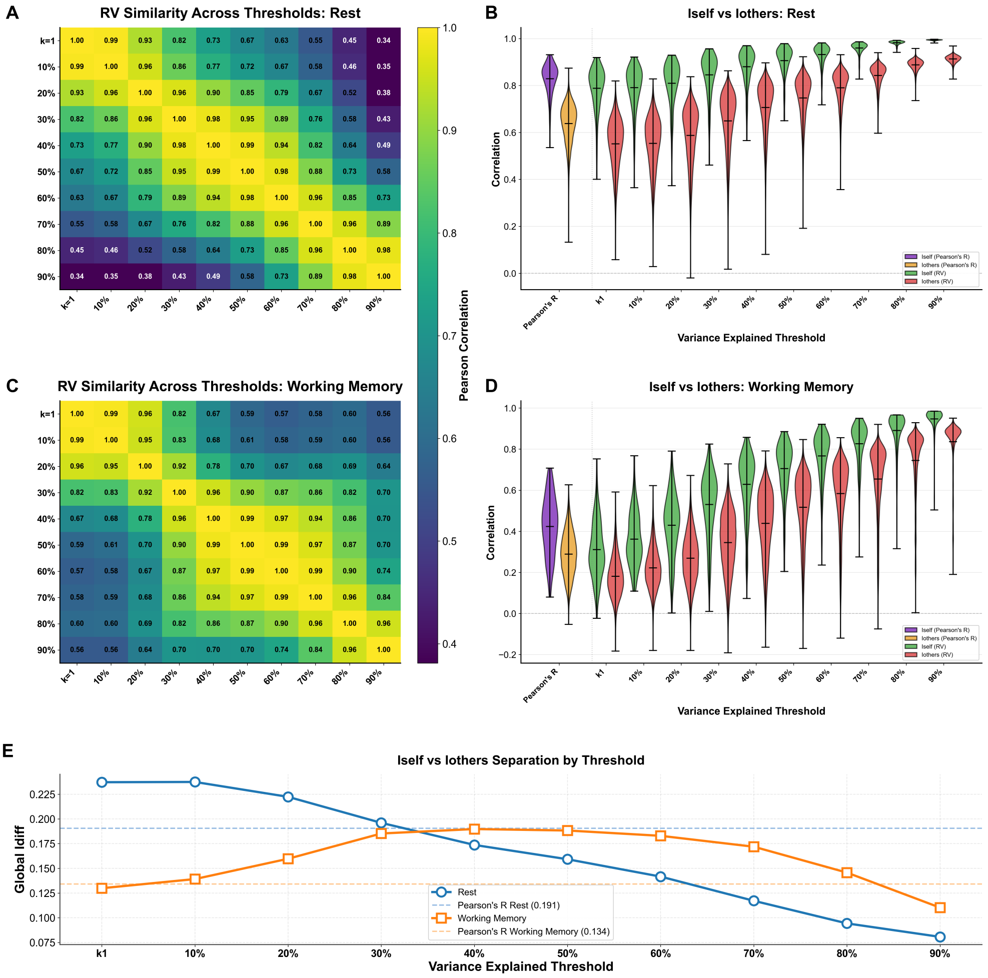


**Supplementary Fig. S24**: **Between-threshold RV similarity and identifiability (Schaefer 200, confirmatory sample).** Same layout as main text Figure 3. A. Rest between-threshold RV similarity. B. Rest $I_{\text{self}}$ vs. $I_{\text{others}}$ violins. C. WM between-threshold RV similarity. D. WM $I_{\text{self}}$ vs. $I_{\text{others}}$ violins. E. $I_{\text{diff}}$ across thresholds (blue: rest, Pearson $I_{\text{diff}} = 0.191$; orange: WM, Pearson $I_{\text{diff}} = 0.134$). The monotonic decline in rest $I_{\text{diff}}$ and inverted-U in WM replicate the Schaefer 100 pattern. The crossover $\tau^*$ at which RV $I_{\text{diff}}$ falls below Pearson $I_{\text{diff}}$ in rest shifts to ≈30–40%.

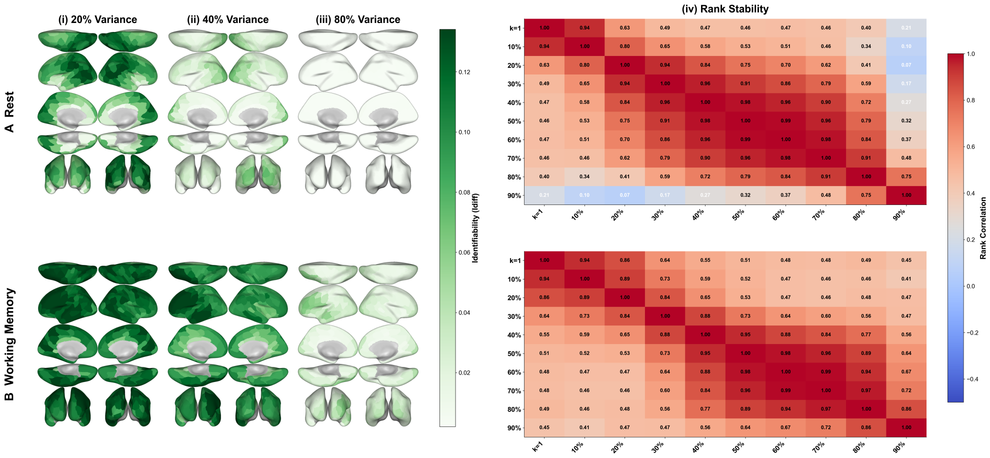


**Supplementary Fig. S25**: **Parcel-level identifiability and rank stability (Schaefer 200, confirmatory sample).** Same layout as main text Figure 5. A. Rest; B. WM. i. $I_{\text{diff}}^{(r)}$ from $\text{RV}_{20}$; ii. $I_{\text{diff}}^{(r)}$ from $\text{RV}_{40}$; iii. $I_{\text{diff}}^{(r)}$ from $\text{RV}_{80}$; iv. rank stability across thresholds. The spatial pattern of identifiability is consistent with Schaefer 100, with visual and somatomotor regions most identifiable at low thresholds.

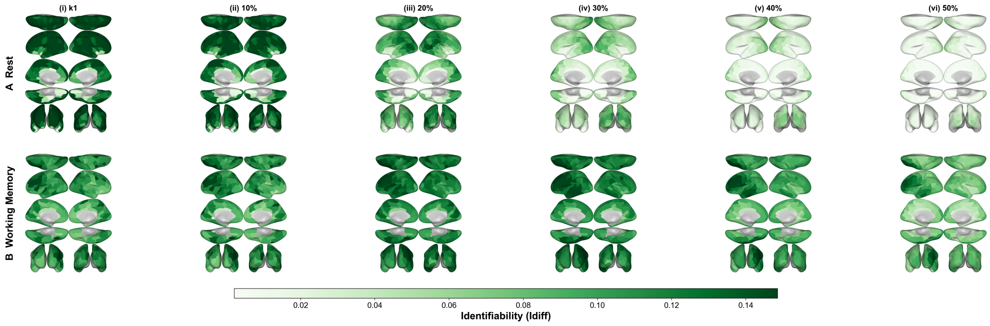


**Supplementary Fig. S26**: **Parcel-level identifiability across low-to-moderate thresholds (Schaefer 200, confirmatory sample).** A. Rest; B. WM. Columns: $I_{\text{diff}}^{(r)}$ from i. $\text{RV}_{k1}$, ii. $\text{RV}_{10}$, iii. $\text{RV}_{20}$, iv. $\text{RV}_{30}$, v. $\text{RV}_{40}$, vi. $\text{RV}_{50}$.

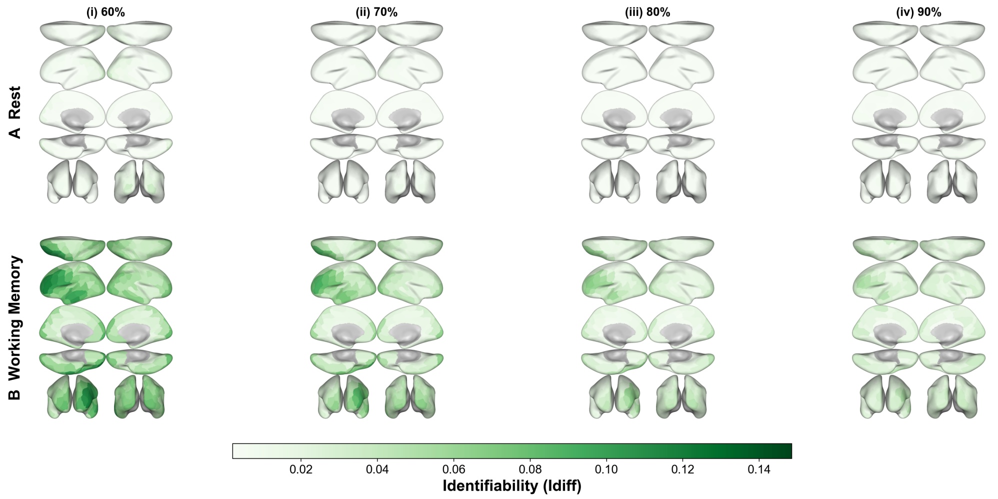


**Supplementary Fig. S27**: **Parcel-level identifiability across high thresholds (Schaefer 200, confirmatory sample).** A. Rest; B. WM. Columns: $I^{(r)}_{\text{diff}}$ from i. $\text{RV}_{60}$, ii. $\text{RV}_{70}$, iii. $\text{RV}_{80}$, iv. $\text{RV}_{90}$.

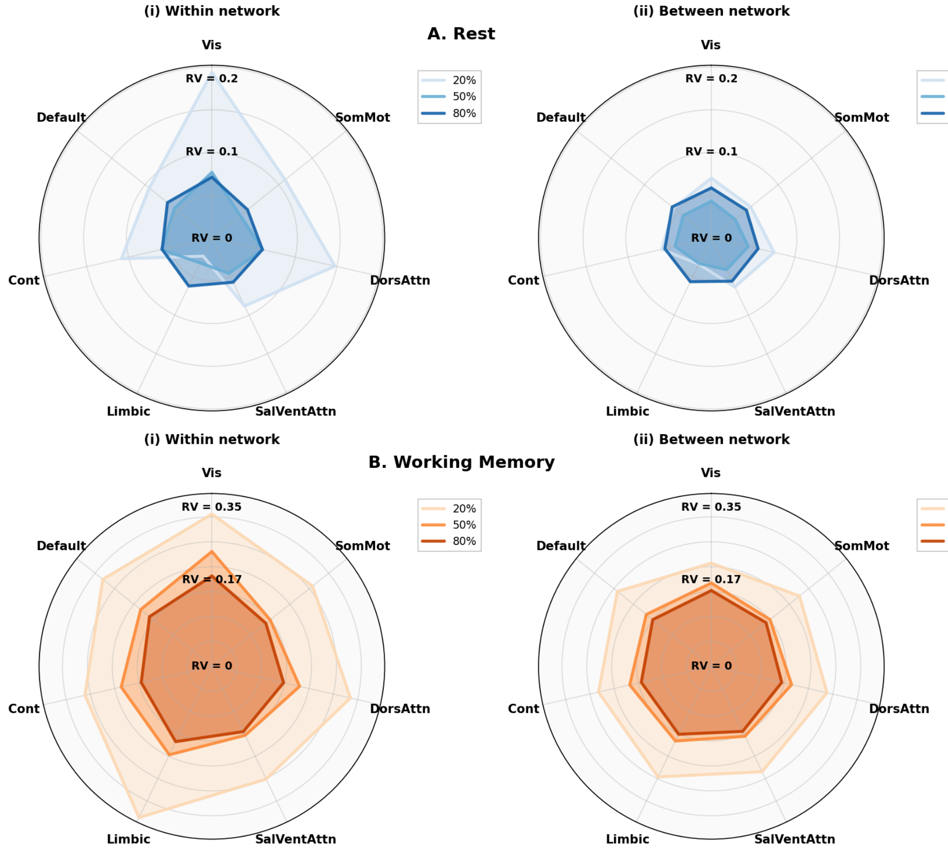


**Supplementary Fig. S28**: **Network-level RV radar plots (Schaefer 200, confirmatory sample).** Same layout as main text Figure 4. A. Rest; B. WM. i. Within-network mean RV; ii. Between-network mean RV. Thresholds: $RV_{20}$, $RV_{50}$, $RV_{80}$. Network segregation at rest (high within, low between at $RV_{20}$) and integration in WM replicate the Schaefer 100 pattern.

#### Schaefer 400 parcellation

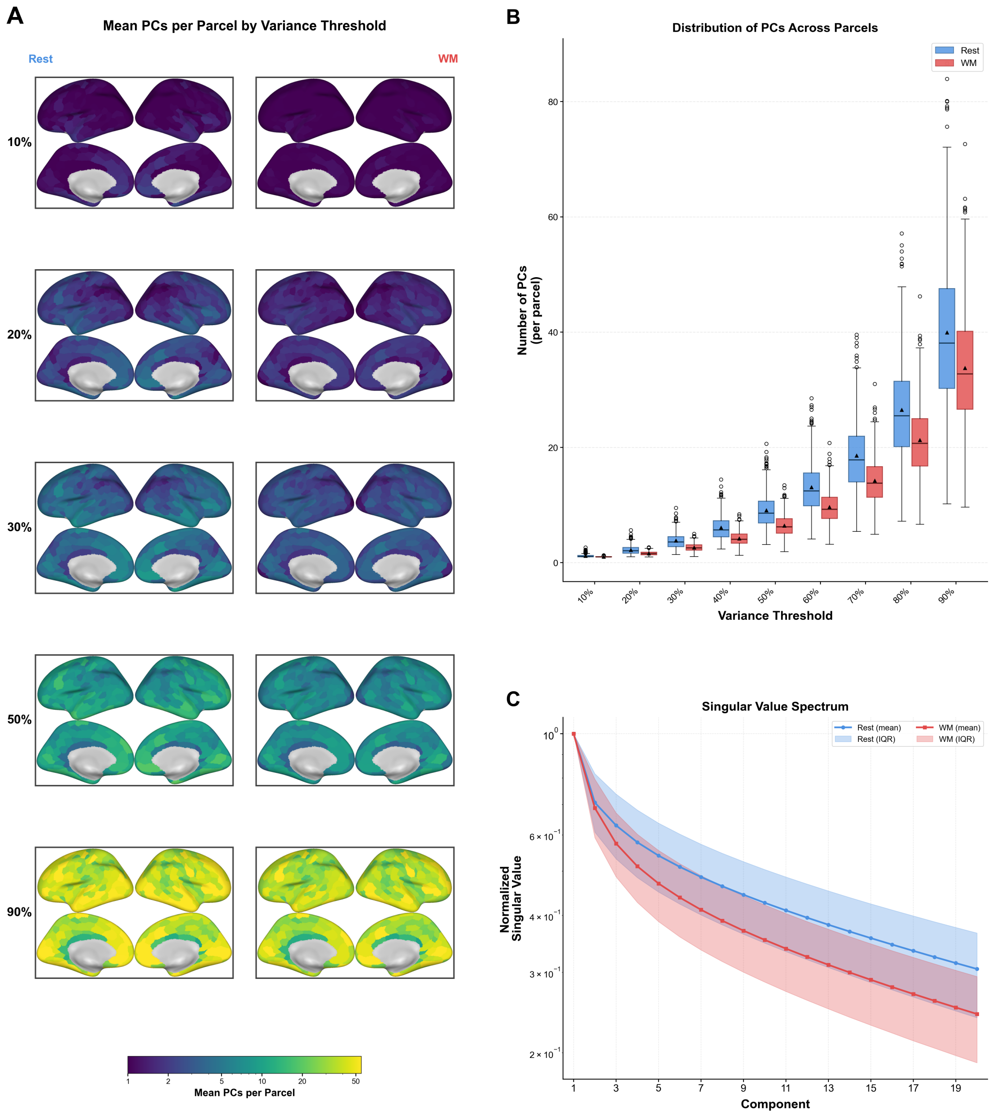


**Supplementary Fig. S29**: **PCs needed to explain within-ROI variance (Schaefer 400, confirmatory sample).** Same layout as main text Figure 2 at the 400-parcel resolution. Absolute PC counts are lower than Schaefer 100 and 200 owing to fewer grayordinates per parcel, but the spatial heterogeneity pattern is preserved.

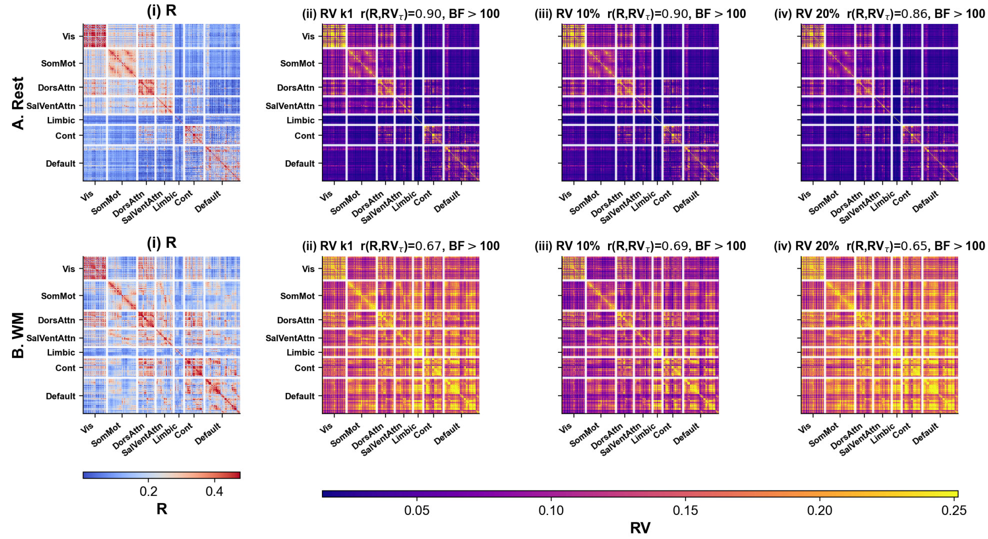


**Supplementary Fig. S30**: **Group FC matrices at low variance thresholds (Schaefer 400, confirmatory sample).** A. Rest; B. WM. i. Pearson's R. RV-based FC at ii. $k_1$, iii. 10%, iv. 20% variance explained. $r(R, \mathrm{RV}_{k1}) = 0.90$ for rest, consistent with Schaefer 100 ($r = 0.92$) and Schaefer 200 ($r = 0.92$).

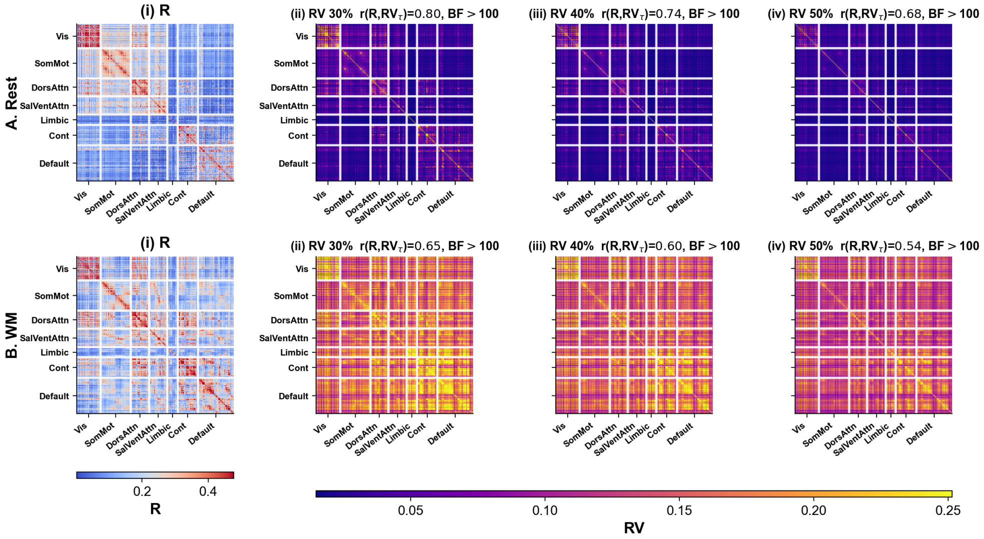


**Supplementary Fig. S31**: **Group FC matrices at moderate variance thresholds (Schaefer 400, confirmatory sample).** A. Rest; B. WM. i. Pearson's R. RV-based FC at ii. 30%, iii. 40%, iv. 50% variance explained. $r(R, \mathrm{RV}_\tau)$ values are the highest of the three parcellations at matched thresholds (e.g., $r(R, \mathrm{RV}_{50}) = 0.68$ vs. 0.57 at Schaefer 200 and 0.43 at Schaefer 100 for rest).

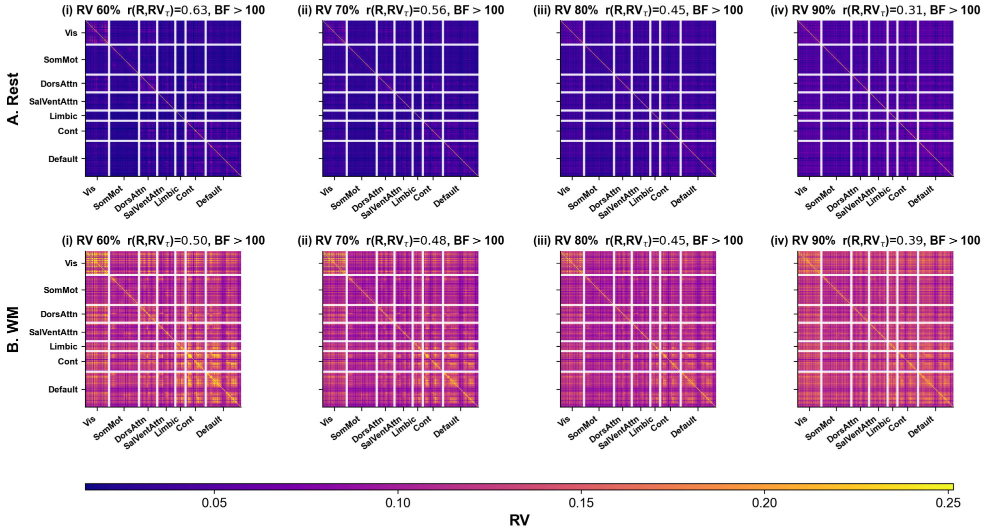


**Supplementary Fig. S32**: **Group FC matrices at high variance thresholds (Schaefer 400, confirmatory sample).** A. Rest; B. WM. RV-based FC at i. 60%, ii. 70%, iii. 80%, iv. 90% variance explained. At $\mathrm{RV}_{80}$, rest $r(R, \mathrm{RV}_{80}) = 0.45$ compared to 0.32 (Schaefer 200) and 0.14 (Schaefer 100), confirming that finer parcellations retain more R-like structure at high thresholds.

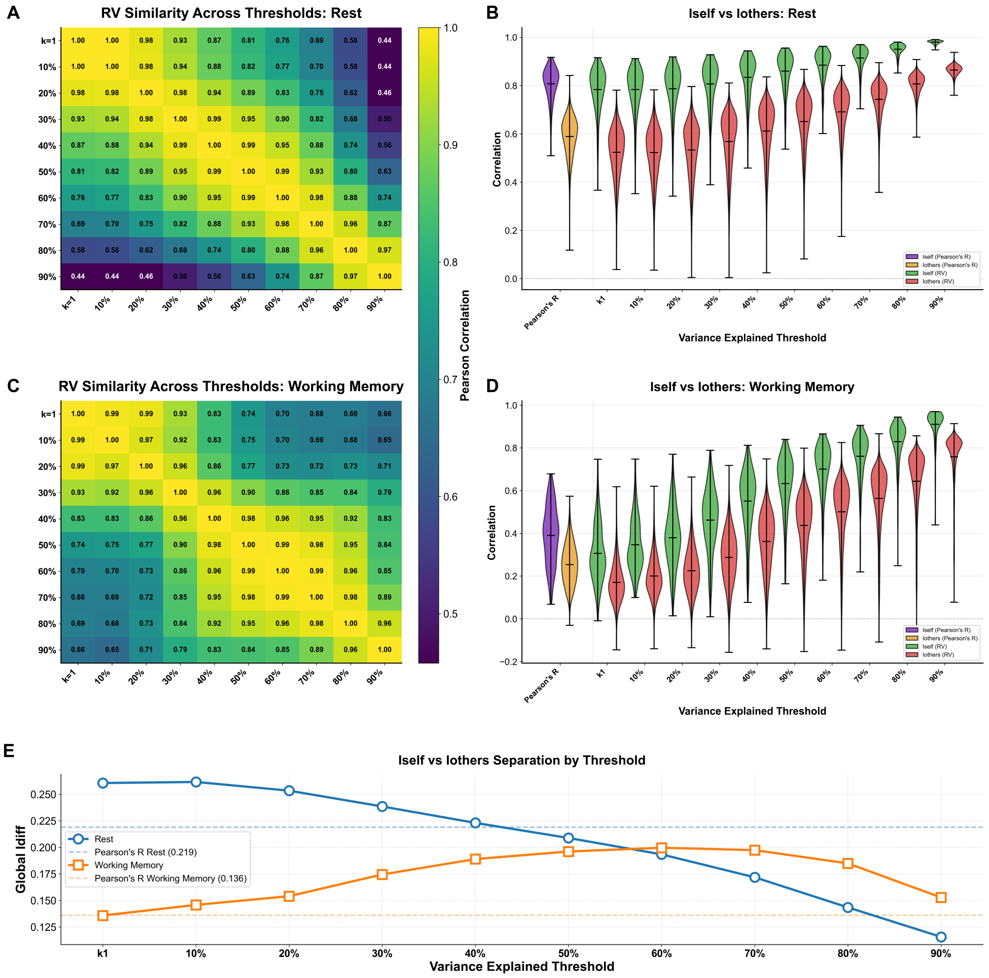


**Supplementary Fig. S33**: **Between-threshold RV similarity and identifiability (Schaefer 400, confirmatory sample).** Same layout as main text Figure 3. E. $I_{\text{diff}}$ across thresholds (blue: rest, Pearson $I_{\text{diff}} = 0.219$; orange: WM, Pearson $I_{\text{diff}} = 0.136$). Peak rest $I_{\text{diff}} \approx 0.26$, the highest of the three parcellations, reflecting increased spatial specificity. The crossover $\tau^*$ shifts to ≈40–50%.

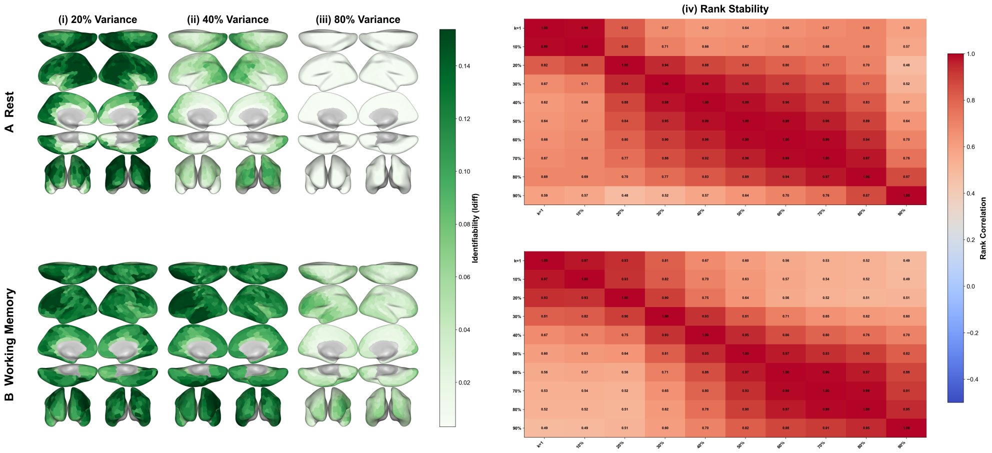


**Supplementary Fig. S34**: **Parcel-level identifiability and rank stability (Schaefer 400, confirmatory sample).** Same layout as main text Figure 5. A. Rest; B. WM. i. $I_{\text{diff}}^{(r)}$ from $\text{RV}_{20}$; ii. $I_{\text{diff}}^{(r)}$ from $\text{RV}_{40}$; iii. $I_{\text{diff}}^{(r)}$ from $\text{RV}_{80}$; iv. rank stability across thresholds. The spatial pattern and three-block rank stability structure are consistent with Schaefer 100 and 200.

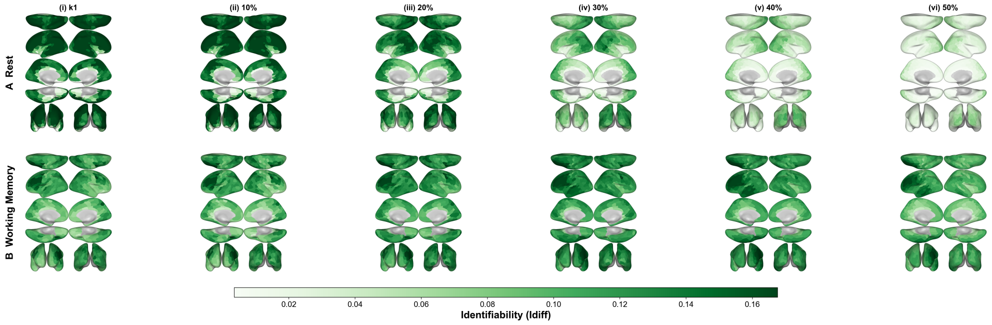


**Supplementary Fig. S35**: **Parcel-level identifiability across low-to-moderate thresholds (Schaefer 400, confirmatory sample).** A. Rest; B. WM. Columns: $I_{\text{diff}}^{(r)}$ from i. $\text{RV}_{k1}$, ii. $\text{RV}_{10}$, iii. $\text{RV}_{20}$, iv. $\text{RV}_{30}$, v. $\text{RV}_{40}$, vi. $\text{RV}_{50}$.

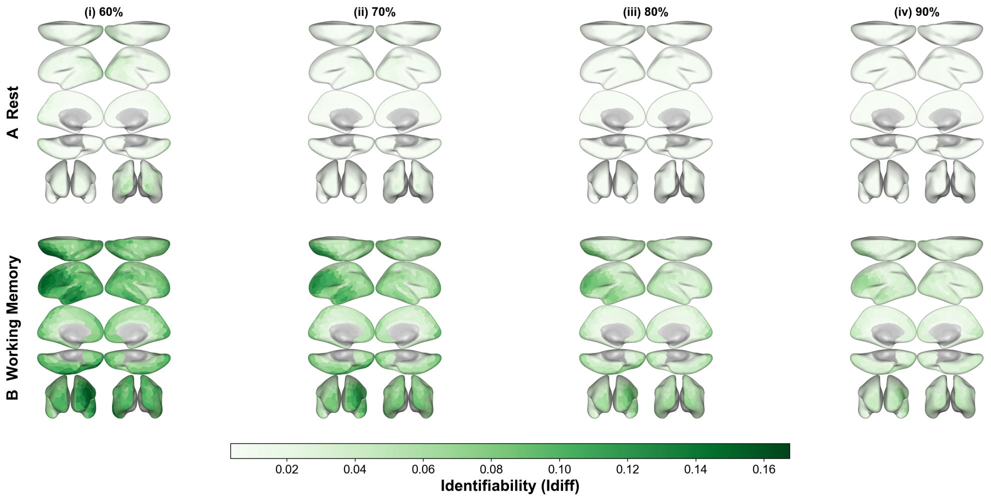


**Supplementary Fig. S36**: **Parcel-level identifiability across high thresholds (Schaefer 400, confirmatory sample).** A. Rest; B. WM. Columns: $I_{\text{diff}}^{(r)}$ from i. $\text{RV}_{60}$, ii. $\text{RV}_{70}$, iii. $\text{RV}_{80}$, iv. $\text{RV}_{90}$.

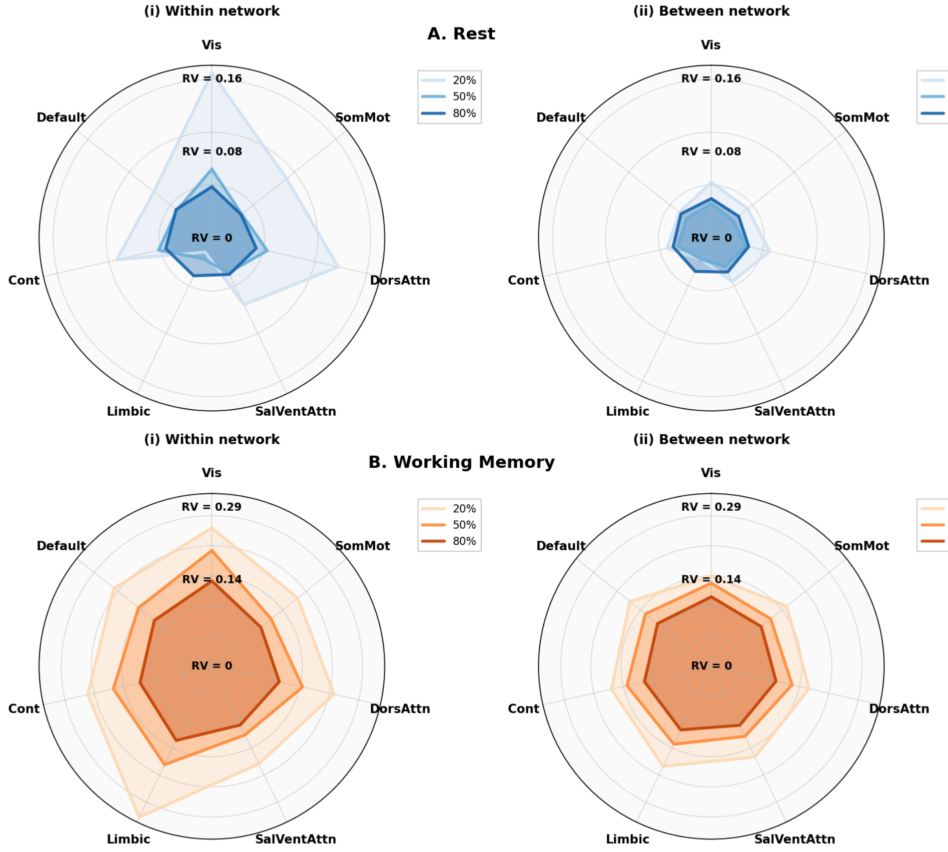


**Supplementary Fig. S37**: **Network-level RV radar plots (Schaefer 400, confirmatory sample).** Same layout as main text Figure 4. A. Rest; B. WM. i. Within-network mean RV; ii. Between-network mean RV. Thresholds: $RV_{20}$, $RV_{50}$, $RV_{80}$. Network segregation at rest and integration in WM are preserved at the finest parcellation.
